# Patient-specific metabolic fluxes reflect molecular subtype and heterogeneity in breast cancer

**DOI:** 10.1101/2025.07.29.667003

**Authors:** Eduardo Ruiz Robles, Rigoberto Rincón-Ballesteros, Sarai Alessandra Chacón Méndez, Francisco J. Alvarez-Padilla, Germán Preciat

## Abstract

Tumor metabolism is dynamic and context-dependent, yet gene expression provides only an indirect view of metabolic activity. We combined patient-specific genome-scale metabolic models with supervised and unsupervised machine learning to infer metabolic fluxes for 1,223 TCGA breast tumor and normal samples under a uniform physiological medium. Flux profiles distinguished tumor from normal tissue under patient-grouped nested cross-validation (ROC-AUC up to 0.989) and predicted receptor status and molecular subtype (e.g., Basal-like versus other tumors, AUC = 0.966), but not stage, nodal status or age. Within tumors, metabolic variation was continuous rather than organized into discrete groups and aligned only partially with clinical classifications. These findings support a view of breast cancer metabolism as a structured but continuous landscape that reflects molecular subtype and receptor status, and provide a reproducible framework to characterize tumor heterogeneity.

## Introduction

Cancer is fundamentally a disease of dysregulated cellular function, yet most current approaches to tumor characterization rely on static molecular measurements that only indirectly reflect cellular behavior [1–3]. While genomic and transcriptomic profiling have transformed cancer classification and enabled advances in precision medicine [1,3], these approaches primarily quantify molecular abundance rather than cellular activity, limiting their ability to capture the processes that ultimately determine tumor phenotypes. Among these processes, metabolism plays a central role by integrating genetic alterations with environmental and microenvironmental constraints, thereby shaping the functional behavior of tumor cells [4,5].

Metabolic reprogramming has long been recognized as a hallmark of cancer, often associated with increased glycolysis and biosynthetic demand [6]. However, accumulating evidence indicates that tumor metabolism is neither uniform nor static, but instead spans a spectrum of context-dependent states shaped by nutrient availability, microenvironmental pressures, and evolutionary dynamics [4,7,8]. This variability suggests that metabolic phenotypes cannot be fully understood through measurements of gene expression alone, but require approaches that capture how metabolic networks function within the limits imposed by cellular and environmental constraints [4,9].

Genome-scale metabolic models (GEMs) provide a mechanistic framework to address this challenge by integrating omics data into network-constrained representations of metabolism [4,5,10,11]. Within this framework, metabolic fluxes, the rates of biochemical reactions, can be inferred as feasible solutions that satisfy stoichiometric, thermodynamic, and gene expression constraints, thereby approximating the set of metabolic states a cell can physically realize. Unlike gene expression, fluxes provide a more direct proxy of cellular activity because they reflect the coordinated execution of metabolic functions under physicochemical constraints [4,9,12]. Under the steady-state assumption, flux distributions describe how metabolites are produced and consumed across the network, thereby influencing energy generation, biosynthesis, and redox balance [10,11], and advances in context-specific model reconstruction and multi-omics integration have further improved the biological relevance of these approaches [12–14].

Recent work has combined GEMs with machine learning to enhance predictive performance and identify clinically relevant biomarkers [12,15–18]. These studies demonstrate the value of integrating mechanistic modeling with data-driven methods, particularly in high-dimensional biological settings [2,15,16]. Notably, prior studies based on gene expression have already demonstrated that metabolic features can stratify patients and predict clinical outcomes [19], further supporting the clinical relevance of metabolism-informed representations. However, in most cases, flux-derived features are used primarily to improve prediction, without explicitly examining the extent to which they encode intrinsic functional organization. As a result, a fundamental question remains open: to what extent do metabolic fluxes capture biologically meaningful information across different scales of tumor organization?

In this study, we address this question by evaluating metabolic fluxes as a quantitative representation of tumor function across multiple biological scales. We develop an integrative framework that combines patient-specific metabolic modeling with supervised and unsupervised learning to systematically assess the informational content of flux distributions [14,20,21]. Using breast cancer as a model system and leveraging large-scale resources such as The Cancer Genome Atlas (TCGA) [22], we show that flux-derived features (i) accurately discriminate tumor from normal tissue, (ii) reveal metabolic heterogeneity across patients, and (iii) contain information predictive of clinically relevant tumor characteristics, such as receptor status and molecular subtype [7,23]. Accordingly, the analyses move from existence to organization to interpretation: whether the flux space carries a reproducible biological signal (tumor versus normal tissue), how metabolic variation is organized among tumors and whether it reproduces clinical classifications, and which clinical characteristics this variation reflects.

Flux-derived features reflect the organization of breast cancer by receptor status and molecular subtype: Basal-like and triple-negative tumors occupy distinct regions of flux space, whereas overall metabolic variation among tumors forms a continuous landscape that only partly follows clinical subtypes [8].

Flux distributions thus link network-level metabolic constraints to clinically observable tumor phenotypes and allow tumor metabolism to be characterized as a spectrum of states [1,24].

## Materials and methods

This study presents an integrative systems biology framework to characterize breast cancer metabolism across multiple functional scales, from tumor discrimination to intratumoral heterogeneity (Fig 1). Patient-specific GEMs were reconstructed from transcriptomic data and used to infer metabolic flux distributions. These fluxes were subsequently analyzed using both supervised and unsupervised learning to assess their ability to discriminate tumor states and reveal intrinsic metabolic heterogeneity.

**Fig 1.**
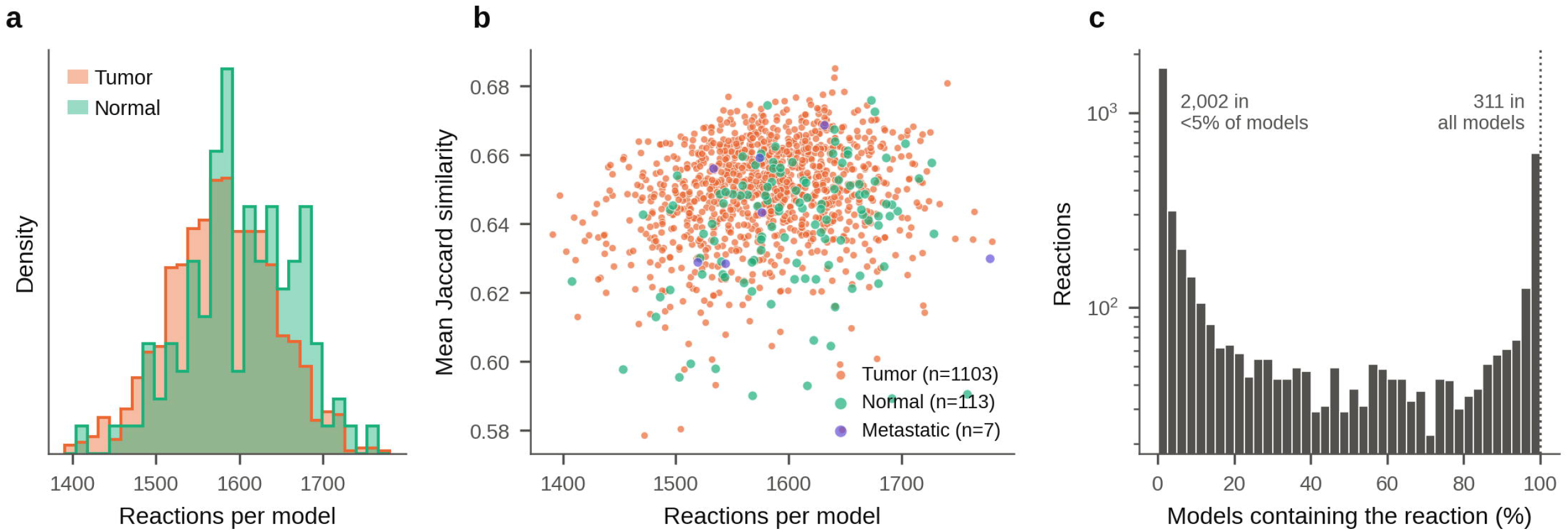
Integrative workflow for multiscale metabolic characterization of breast cancer. **a**) Clinical and transcriptomic data were obtained from the TCGA-BRCA project [22]. b) A curated set of breast cancer–associated metabolic genes, reactions, and metabolites was incorporated as prior knowledge to guide model reconstruction. c) Patient-specific genome-scale metabolic models were reconstructed by integrating transcriptomic data with curated constraints. d) Metabolic fluxes were estimated under a uniform medium using complementary constraint-based optimization strategies. e) Supervised learning was applied to assess tumor–normal discrimination. f) Unsupervised clustering explored metabolic heterogeneity among tumors. g) Metabolic profiles were integrated with clinical variables for multiscale interpretation.

### Data Acquisition and Cohort Definition

Clinical and gene expression data from the TCGA-BRCA project were obtained for both tumor and normal breast tissue samples from the GDC Data Portal and the UCSC Xena platform [22]. Clinical data included 17 variables such as age at diagnosis, sex, race, and tumor characteristics. The gene expression matrix comprised 60,660 Ensembl gene identifiers per sample; after removing 44 duplicated pseudoautosomal (PAR_Y) identifiers, 60,616 genes remained. Expression (STAR-aligned gene counts, GENCODE v36 annotation), clinical and survival tables were obtained from the GDC TCGA-BRCA project through the UCSC Xena GDC hub (data files released in 2024). Expression values were distributed as log-transformed FPKM (FPKM_log_, Eq 1) and were back-transformed to FPKM before model reconstruction:

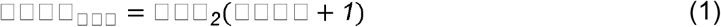

No sample selection was applied: all 1,226 TCGA-BRCA samples with RNA-seq profiles (primary tumors, solid normal tissue, and metastases; 1,095 patients) were used to reconstruct one patient-specific model per sample. Models whose reconstruction failed quality control (n = 3; empty model files) were excluded, leaving 1,223 models (1,103 primary tumors, 113 normal tissues and 7 metastases). All 1,216 primary tumor and normal samples (1,092 patients) were used for the supervised analysis; metastases were excluded because of their small number. For unsupervised and clinical analyses, one primary tumor sample per patient was retained (1,092 tumors). Sample types were assigned from the TCGA barcode (01, primary tumor; 11, solid normal tissue; 06, metastasis), and the first 12 characters of the barcode (TCGA-XX-XXXX) identified the patient for all patient-grouped splits. For patients with more than one primary tumor sample (11 patients), the first vial in barcode order was retained. Sample identifiers were replaced by sequential codes in all Supporting Information.

### Construction of Patient-Specific Metabolic Models

Patient-specific GEMs were reconstructed from the generic human metabolic network Recon3D [11], which includes 10,600 reactions, 5,835 metabolites, and 2,248 genes. Transcriptomic profiles were mapped to Recon3D gene–protein–reaction (GPR) associations using the XomicsToModel framework with the ThermoKernel algorithm [13]; genes with an expression of at least e² FPKM (≈7.4 FPKM; transcriptomic threshold of 2 on the natural logarithmic scale) were considered active (the XomicsToModel default, which retains moderately and highly expressed genes). All reconstruction settings and quality-control criteria are listed in S2 Table. Gene expression levels obtained from bulk RNA-seq data of 1,106 primary tumor samples (1,103 after quality control), 113 solid normal tissue samples, and 7 metastatic samples were used to prioritize the inclusion of reactions supported by transcriptomic evidence while preserving network connectivity and thermodynamic consistency, enabling the reconstruction of patient-specific metabolic networks.

To improve biological relevance, a curated set of cancer-associated metabolic genes (S1 File) was incorporated as additional constraints during reconstruction. This step guided model extraction toward disease-relevant metabolic configurations while preserving network consistency. The same curated constraints were applied to every tumor and normal model, so they cannot by themselves create tumor–normal or subtype differences.

### Estimation of Metabolic Flux Distributions

The space of feasible metabolic flux distributions is inherently non-unique, as multiple flux configurations can satisfy stoichiometric and physicochemical constraints. Rather than representing a single metabolic state, this solution space reflects a spectrum of possible metabolic strategies available to tumor cells.

To ensure that differences between models reflect transcriptomic variation rather than differences in nutrient availability, all models were simulated under the same physiological medium (S4 Table). Uptake was allowed only for the components of Human Plasma-Like Medium (HPLM) [25], serum lipids (palmitate, oleate, linoleate, linolenate, arachidonate and cholesterol), inorganic species, and a small set of biomass precursors that context-specific models cannot synthesize; every other exchange reaction was closed to uptake. Uptake bounds were scaled to the HPLM concentration of each component relative to glucose (5 mM, bound = 1). Models that could not reach the minimal biomass flux under this medium received the smallest additional uptake required, which was recorded for each model and tested as a potential confounder. All optimization problems were solved using deterministic solvers (Gurobi) to avoid variability due to numerical instability.

Because metabolic networks admit multiple feasible flux configurations, we employed complementary optimization strategies to sample distinct regions of the solution space rather than relying on a single optimal solution.

To systematically explore this space, multiple constraint-based optimization approaches were applied:

● Flux Balance Analysis (FBA): Maximizes biomass production to approximate proliferative metabolic states and cellular growth potential under steady-state constraints [26].
● Parsimonious FBA (pFBA): Minimizes the sum of metabolic fluxes while maintaining the optimal growth rate, yielding a flux distribution that approximates minimal enzyme usage while preserving maximal growth (99% of the FBA optimum) [27].
● Weighted L1 minimization (L1w): Minimizes the expression-weighted sum of absolute fluxes (Eq 2) while maintaining 99% of the FBA optimum. The weight of each reaction decreases with its expression (Eq 3), so that weakly expressed reactions are more costly and flux is preferentially routed through expressed reactions [28,29]. Reaction expression was obtained by mapping gene FPKM values through the gene–protein–reaction rule, log-transformed with the natural logarithm and min–max scaled to [0, 1] within each model (Eq 4), where *v*_j_ is the flux of reaction *j* and the minimum and maximum are taken over the reactions of the model with expression data. Reactions without gene association received the median weight.

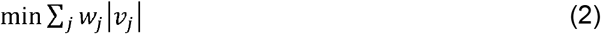

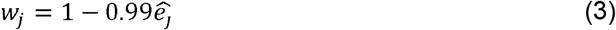

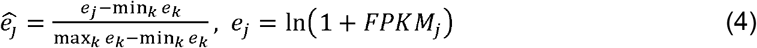
● Multi-objective optimization: Captures heterogeneous metabolic phenotypes by simultaneously considering competing biological objectives (biomass production, ATP generation and carbon uptake), using a 10 × 10 grid of biomass and ATP levels with minimal carbon uptake at each point (S3 Fig) [30].

Unweighted and weighted L2-norm minimizations were also computed. They yielded nearly identical flux profiles (median correlation 0.99) that were also strongly correlated with pFBA and L1w (median correlation ≥ 0.85) and carried flux through almost the whole network; being largely redundant with the other solutions, they are reported only in S1 Fig and were not used in the main analyses.

Following flux estimation, secondary metabolic metrics were calculated from the resulting flux distributions, including ATP consumption, carbon consumption, enzymatic abundance, and other derived metabolic indicators. Subsystem activity was defined as the mean absolute flux of the subsystem reactions present in each model. Fluxes were used as signed values; a flux was considered nonzero if |*v*| > 10^−9^; reactions absent from a model were assigned zero flux; and features nonzero in fewer than 5% of samples were discarded. Definitions of all metrics are given in S2 Table.

Each strategy generated a distinct representation of the metabolic phenotype of the tumors, allowing us to assess the ability of different optimization frameworks to capture inter-tumoral metabolic heterogeneity. The three solutions were treated as complementary representations of the same patient-specific metabolic state. The resulting metabolic representations (FBA, pFBA, L1w, and their combination) were subsequently subjected to dimensionality reduction, separately for each strategy and for the combined set, to identify metabolic subgroups among patients.

### Supervised Learning: Validation of Metabolic Signal

To evaluate whether metabolic fluxes contain discriminative biological information, supervised learning was applied to classify tumor versus normal samples. The dataset comprised all 1,216 primary tumor and normal samples (1,103 tumor and 113 normal; 90.7% and 9.3%), reflecting the class imbalance of the TCGA-BRCA cohort. Input features were the 2,863 reaction fluxes and derived metrics of the FBA, pFBA and L1w solutions (not the pathway summaries shown in Fig 3). Because several patients contributed both a tumor and a normal sample, all partitions were grouped by patient so that no patient appeared in both training and evaluation sets. Performance was estimated by nested cross-validation: an outer patient-grouped, stratified 5-fold split (StratifiedGroupKFold), repeated with three random seeds (42, 123 and 100), and an inner 3-fold split used for randomized hyperparameter search. In addition, a patient-grouped, stratified 70/30 split was used as an independent test set (364 samples) that was not used for model selection. Class imbalance was addressed with balanced class weights in all algorithms that support them (logistic regression, SVM, random forest, gradient boosting and decision tree), and imbalance-robust metrics (specificity, balanced accuracy and Matthews correlation coefficient) were reported together with a majority-class baseline classifier.

Feature selection was performed at the reaction level: for each reaction flux or derived metric, a two-sided, unpaired Mann–Whitney U test compared tumor and normal samples of the training data. Statistical significance was determined using the Benjamini–Hochberg correction (FDR ≤ 0.05) to control the false discovery rate [31,32]. Feature selection was implemented as a step of the classification pipeline, so it was repeated within every training fold of the nested cross-validation and never used the evaluation samples, avoiding information leakage. Selected features were standardized prior to model training. The number of selected features varied across folds depending on statistical significance after FDR correction.

Eight classification algorithms were benchmarked to test whether the tumor–normal signal is robust across model classes:

● Gradient Boosting (histogram-based gradient boosting, scikit-learn).
● Random Forest.
● K-Nearest Neighbors (KNN).
● Support Vector Machines (SVM).
● Logistic Regression.
● Multi-Layer Perceptron (MLP).
● Decision Tree.
● Naive Bayes.

Hyperparameters were optimized for each algorithm in the inner cross-validation loop by randomized search (six candidate settings per algorithm and fold) maximizing ROC-AUC; search spaces and the selected values are provided in S3 Table. A majority-class (dummy) classifier was evaluated with the same procedure as a reference. Classifiers were implemented in scikit-learn, and all analyses were executed using fixed random seeds when stochastic procedures were involved.

### Unsupervised Learning: Exploration of Metabolic Heterogeneity

Following validation of the metabolic signal, unsupervised learning was used to explore metabolic heterogeneity among primary tumors. Flux-derived metrics were transformed with a signed logarithm, standardized, reduced to 50 principal components, and embedded in two or three dimensions with Principal Component Analysis (PCA) and Uniform Manifold Approximation and Projection (UMAP; 15 or 30 neighbors, minimum distance 0.1, Euclidean metric, three random seeds).

Clustering was performed using multiple algorithmic families representing complementary mathematical principles:

● Partition-based methods.
● Hierarchical clustering.
● Density-based clustering.
● Probabilistic models.

To ensure robustness, parameter sweeps and multiple random initializations were performed (k-means, agglomerative clustering, Gaussian and Bayesian Gaussian mixtures, DBSCAN, HDBSCAN and mean shift), and solutions whose smallest cluster contained less than 2% of the tumors were discarded. Clustering results were compared across flux distributions derived from different optimization strategies, allowing assessment of whether observed phenotypes are consistent across alternative metabolic states. Each solution was evaluated with silhouette, Davies–Bouldin and Calinski–Harabasz indices, computed both in the embedding and in the original feature space. Partitions associated with missing clinical metadata, tissue source site or medium supplementation (Cramér’s V ≥ 0.3 or p < 0.001) were discarded, and differences in model size were reported. The final partition was chosen by a pre-defined protocol: stable across UMAP seeds (mean adjusted Rand index, ARI ≥ 0.5), not based on FBA metrics alone (FBA optima are not unique), and with the highest silhouette in the original feature space.

This approach enables the exploration of metabolic structure independent of predefined clinical labels.

### Integration with Clinical Variables

Three datasets were retrieved from the GDC Data Portal and the UCSC Xena platform, comprising more than 85 variables describing clinical, demographic, pathological, therapeutic, and survival characteristics of the patients. The dataset was analyzed using statistical methods and data visualization techniques to identify variables that could reflect metabolic states and tumor prognosis. Sixteen variables were excluded because they contained more than 97% missing values and were primarily related to sample annotations, institutional information, and identification codes. In addition, new features were derived from existing clinical variables to enrich the dataset. Following data preprocessing, several unsupervised learning models were trained using different combinations of variables. Since clustering algorithms group patients according to the characteristics included in the analysis, the selection of variables has a direct impact on the resulting patient stratification [33]. Therefore, we aimed to identify clinical variables that best captured tumor metabolic behavior, facilitating the comparison and association of patient groups with the metabolic states inferred from the genome-scale metabolic models (GEMs). A refined subset of biologically relevant variables (S1 Table) was then used to cluster the 1,069 tumors with clinical records from clinical data alone; missing categories were encoded as a separate level, and PAM50 (a transcriptomic classifier) was excluded to avoid circularity. To evaluate clinical relevance, metabolic clusters were compared with clinical and pathological variables, including receptor status (ER, PR, HER2), PAM50 subtype, stage, histology and ancestry, using χ² tests with Cramér’s V and Benjamini–Hochberg correction. Overall survival (time from diagnosis to death or last follow-up) was compared with Kaplan–Meier estimates, the log-rank test, and an exploratory Cox model adjusted for age, stage and PAM50 subtype. All clinical and metabolic features used can be found in supplementary material (S1 and S2 Tables).

Agreement between metabolic and clinical groupings was quantified using the ARI, computed on the tumors common to both partitions and compared with a permutation null (1,000 label shuffles) [34].

To quantify which clinical characteristics can be predicted from metabolic fluxes independently of any clustering, a regularized logistic regression (with the same within-fold feature selection) was trained to predict each clinical variable from flux features, and its ROC-AUC was estimated by stratified 5-fold cross-validation repeated with three seeds (balanced class weights; endpoint definitions and class counts in S6 Table). Differences in flux-derived pathway activity among PAM50 subtypes were tested with the Kruskal–Wallis test (FDR ≤ 0.05), and effect sizes between groups were quantified with Cliff’s delta.

### Evaluation Metrics and Model Selection

Supervised model performance was evaluated using:

● Accuracy.
● Precision.
● Sensitivity (recall), specificity and balanced accuracy.
● F1-score and Matthews correlation coefficient (MCC).
● ROC-AUC.

To assess robustness and avoid overfitting, additional metrics were included:

● Generalization index, capturing performance differences between training and validation sets.
● Stability index, measuring variability across cross-validation folds.

A multi-criteria score summed the 11 metrics of Fig 7 with equal weights (all scaled to [0, 1]; MCC truncated at 0; fit time min–max scaled and inverted) to rank the algorithms.

For unsupervised analyses, evaluation prioritized:

● Cluster separation (silhouette, Davies–Bouldin and Calinski–Harabasz indices) and consistency across methods.
● Concordance with clinical variables (adjusted Rand index and adjusted mutual information).

The complete set of evaluation metrics is summarized in Table 1.

**Table 1.**
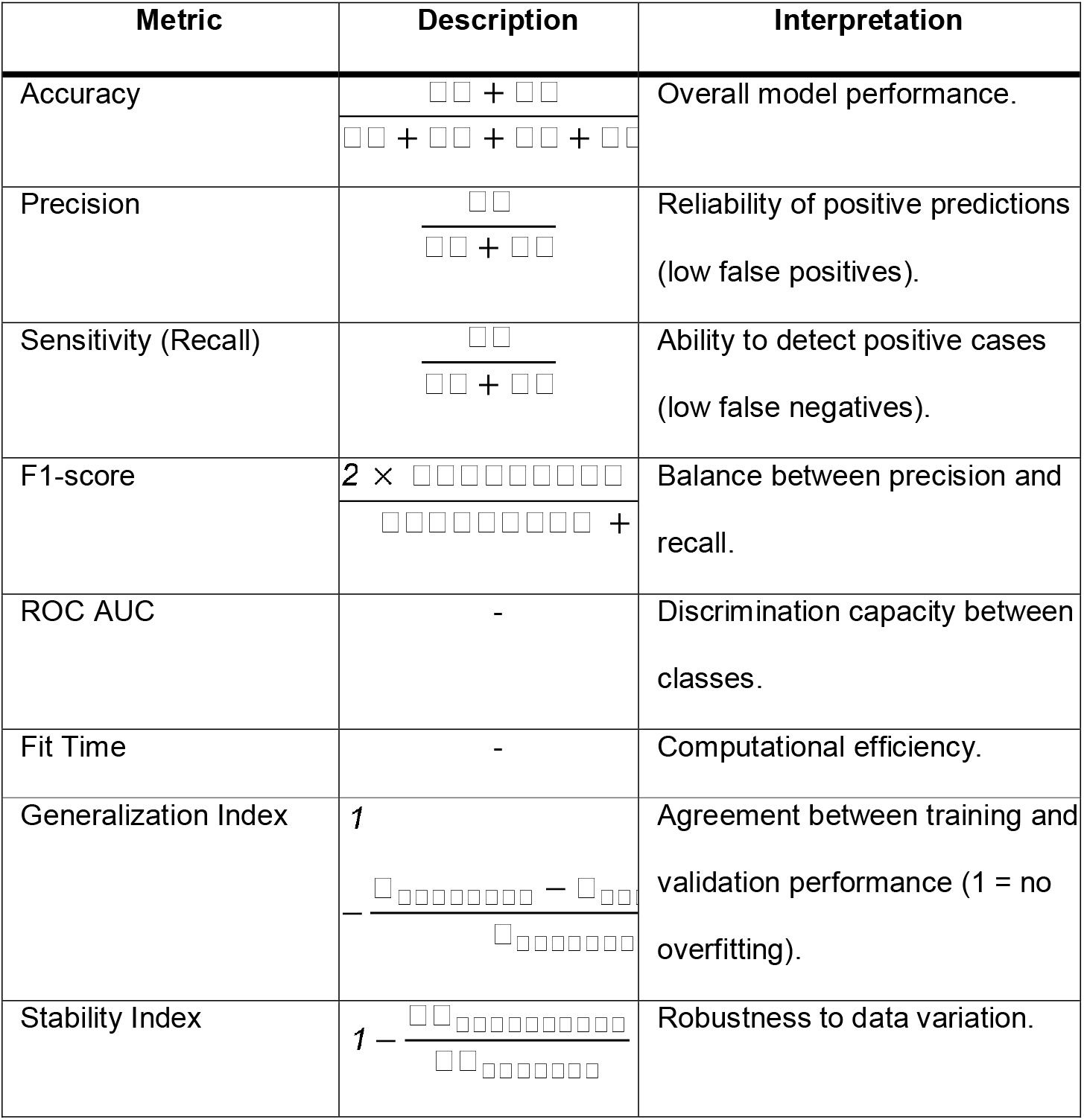
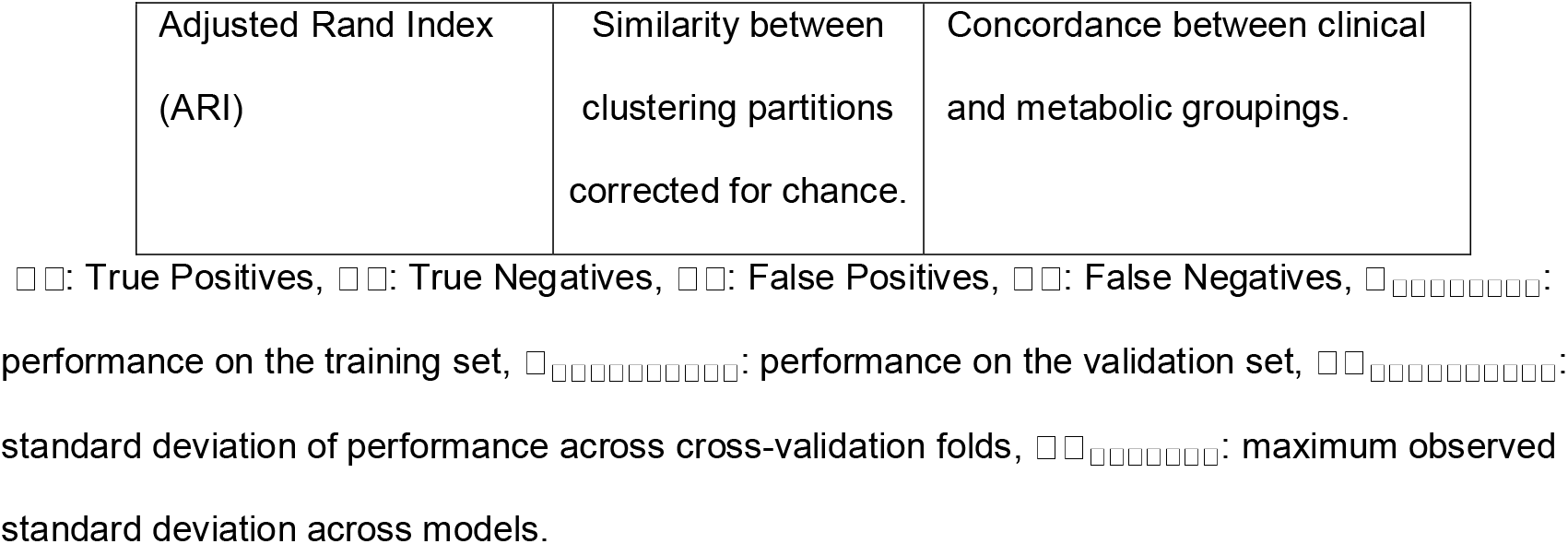
Metrics used for model evaluation and validation.

Together, this framework enables a systematic evaluation of metabolic flux-derived features for both classification and the exploration of metabolic heterogeneity, bridging mechanistic modeling with clinically interpretable tumor characterization.

### Computational Environment

All analyses were implemented using a structured and reproducible pipeline integrating MATLAB (COBRA Toolbox v3.4 [10]), Gurobi for mathematical optimization and Python 3 (scikit-learn, NumPy, SciPy, pandas, Matplotlib). A single MATLAB script computes all flux solutions, derived metrics and quality-control indicators for every model, and a Python notebook generates all statistics, tables and figures reported here. Model reconstruction, flux estimation, and downstream analyses were executed through version-controlled scripts available in the public repository.

Key parameters were fixed across analyses unless otherwise specified:

● Cross-validation: nested, patient-grouped stratified 5-fold (outer) and 3-fold (inner) with three fixed random seeds (42, 123, 100) to assess reproducibility across replicates.
● Feature selection threshold: FDR ≤ 0.05.
● Clustering: each algorithm was run over a parameter grid on PCA and UMAP embeddings computed with three random seeds.

The complete workflow, including data preprocessing, model reconstruction, flux estimation, and statistical analysis, can be executed end-to-end using the provided scripts.

## Results

### Construction and variability of patient-specific metabolic models

The 1,223 patient-specific GEMs were similar in size, with an average of 1,582 ± 64 reactions, 968 metabolites, and 919 genes (Fig 2a).

**Fig 2.**
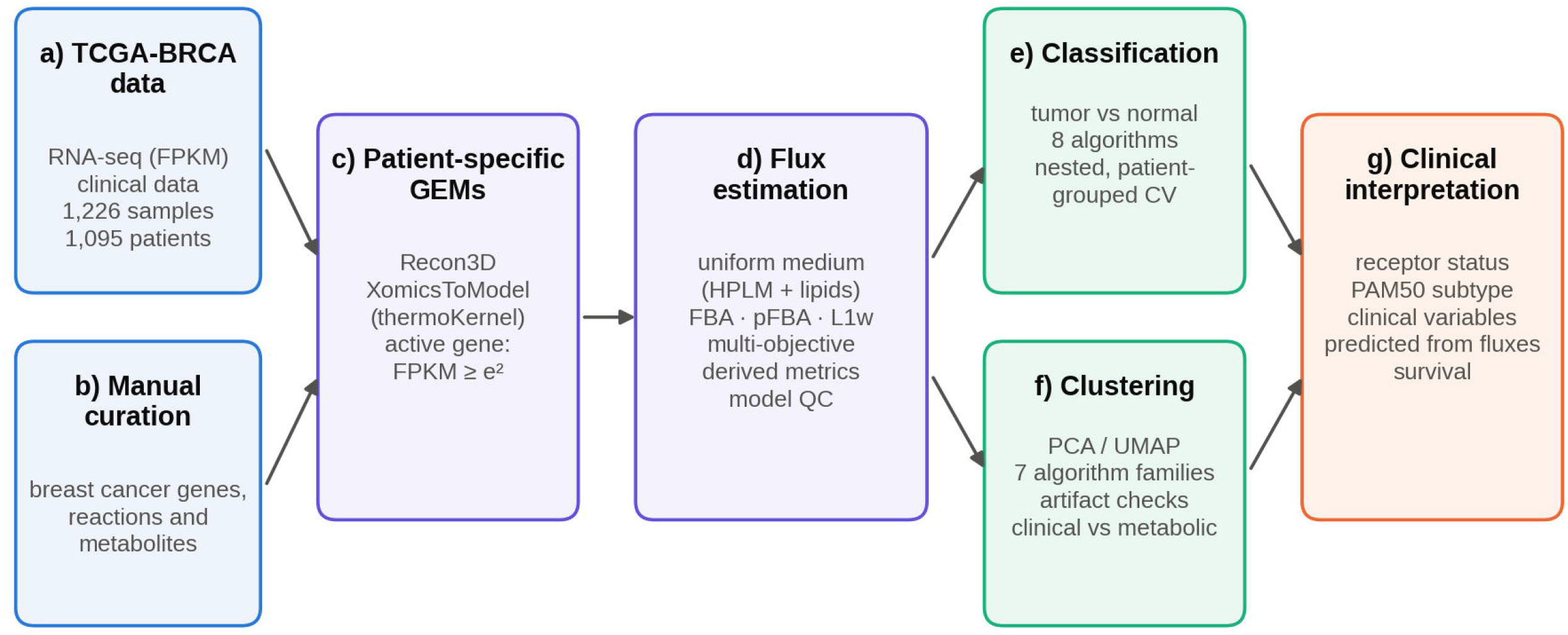
Patient-specific metabolic models show heterogeneous reaction content and structural similarity. (a) Distribution of the number of reactions per model in tumor and normal tissue. (b) Number of reactions versus the mean Jaccard similarity of each model to all other models, colored by tissue type (1,223 GEMs). (c) Fraction of models that contain each reaction; 311 reactions are present in all models and 2,002 in fewer than 5%.

**Fig 3.**
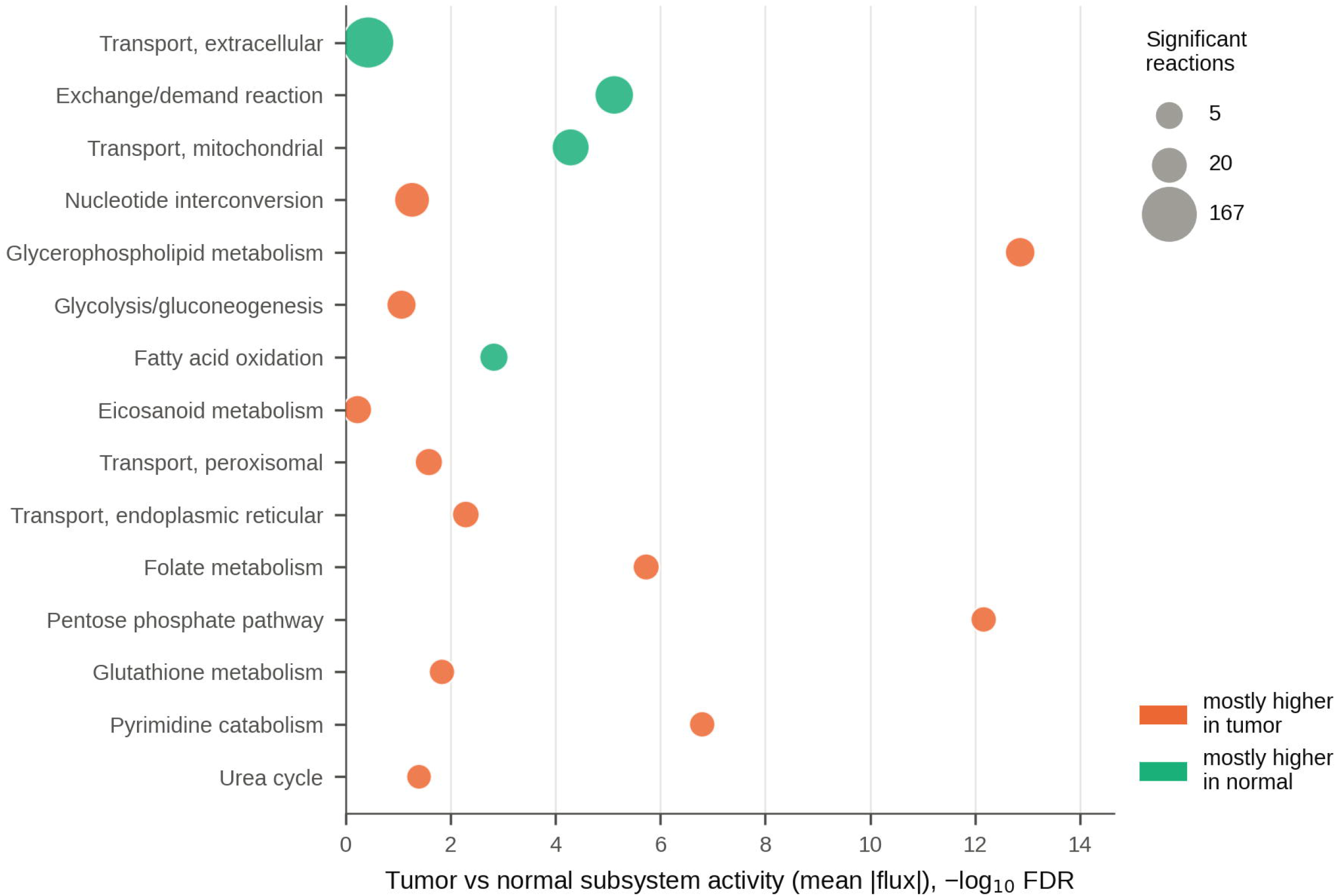
Significantly differentiable metabolic pathways. The 15 subsystems with the largest number of reactions whose flux differs between tumor and normal samples (Mann–Whitney U test, FDR ≤ 0.05). Point size represents the number of significant reactions; the x-axis shows the significance of the difference in mean absolute subsystem flux; color indicates whether most significant reactions were higher in tumor or in normal tissue. Reaction– and gene-level results are given in S5 Table.

Reaction content, however, varied considerably between models. Of the 4,713 reactions present in at least one model, only 311 were shared by all models and 2,002 were present in fewer than 5% of them (Fig 2c), indicating a small common core and a large patient-specific component.

Mean pairwise reaction-content similarity (Jaccard index) was 0.649 ± 0.027 (Fig 2b), was similar when exchange, demand, sink and transport reactions were excluded (0.659), and was higher within than between tissue types (tumor–tumor 0.651, normal–normal 0.671, tumor–normal 0.636). Thus, models of similar size (Fig 2a) differ in which reactions they contain.

These differences describe structural heterogeneity of the reconstructions, not differences in metabolic activity. We therefore asked whether this structural variability translated into differences in metabolic flux.

### Metabolic fluxes capture tumor versus normal differences

To evaluate whether metabolic flux distributions reflect systematic metabolic differences between tumor and normal tissue, each signed reaction flux of the FBA, pFBA and L1w solutions was compared between groups (Mann–Whitney U test, FDR ≤ 0.05; each reaction counted once, with its most significant solution). Of 1,504 reactions carrying flux, 526 differed significantly (284 higher in tumors), and these reactions were summarized by subsystem (Fig 3). The individual reactions, their effect sizes and the associated genes are listed in S5 Table.

The largest numbers of altered reactions were found in extracellular and mitochondrial transport and in exchange reactions, mostly with higher flux in normal tissue, whereas altered reactions in nucleotide interconversion, glycerophospholipid metabolism, glycolysis, the pentose phosphate pathway, folate metabolism and pyrimidine catabolism were mostly higher in tumors; glycerophospholipid metabolism and the pentose phosphate pathway showed the strongest subsystem-level differences. These pathways reflect core functional demands of tumor cells, including membrane synthesis and the production of NADPH and nucleotides for proliferation [6–8].

These differences are not limited to individual reactions but instead reflect coordinated shifts across entire pathways. This suggests that tumor metabolism operates through integrated functional programs rather than isolated pathway alterations.

To validate whether metabolic fluxes contain discriminative biological information, supervised learning models were trained to distinguish tumors from normal samples, using reaction-level fluxes and derived metrics as features (the pathway summaries of Fig 3 are descriptive and were not used as model inputs).

All models achieved high predictive performance under nested, patient-grouped cross-validation, with Gradient Boosting reaching the highest ROC-AUC (0.989) followed by Random Forest (0.985) and SVM (0.981; Table 2). Because 90.7% of samples are tumors, accuracy alone is uninformative (the majority-class baseline reaches 0.907); specificity, balanced accuracy and MCC show that the best models also recognized most normal samples (balanced accuracy 0.818–0.918). Excluding the 37 models that required medium supplementation (36 tumors and 1 normal sample) from the evaluation changed the out-of-fold ROC-AUC of every algorithm by at most 0.007, and a classifier using only reconstruction variables (model size, number of active reactions, supplementation) reached a ROC-AUC of 0.785, well below that of flux-based models (S3 Table).

**Table 2.** Nested cross-validation performance for tumor versus normal classification. Mean over 15 outer folds (5 patient-grouped folds × 3 seeds); standard deviations are given in S3 Table. LR, logistic regression; SVM, support vector machine; KNN, k-nearest neighbors; RF, random forest; GB, gradient boosting; MLP, multi-layer perceptron; DT, decision tree; NB, naive Bayes; Base, majority-class baseline.

| **Metric** | **GB** | **RF** | **SVM** | **LR** | **KNN** | **MLP** | **DT** | **NB** | **Base** |
|---|---|---|---|---|---|---|---|---|---|
| Accuracy | 0.975 | 0.961 | 0.967 | 0.962 | 0.948 | 0.955 | 0.872 | 0.894 | 0.907 |
| Precision | 0.982 | 0.965 | 0.974 | 0.986 | 0.951 | 0.978 | 0.971 | 0.964 | 0.907 |
| Sensitivity | 0.991 | 0.993 | 0.990 | 0.972 | 0.995 | 0.972 | 0.885 | 0.917 | 1.000 |
| Specificity | 0.823 | 0.643 | 0.744 | 0.864 | 0.489 | 0.788 | 0.743 | 0.661 | 0.000 |
| F1-score | 0.986 | 0.979 | 0.982 | 0.979 | 0.972 | 0.975 | 0.926 | 0.940 | 0.951 |
| Balanced accuracy | 0.907 | 0.818 | 0.867 | 0.918 | 0.742 | 0.880 | 0.814 | 0.789 | 0.500 |
| MCC | 0.850 | 0.746 | 0.796 | 0.790 | 0.630 | 0.745 | 0.487 | 0.491 | 0.000 |
| ROC-AUC | 0.989 | 0.985 | 0.981 | 0.969 | 0.959 | 0.946 | 0.837 | 0.809 | 0.500 |

Decision trees and naive Bayes were included as interpretable or simple baselines and showed the lowest performance (ROC-AUC 0.837 and 0.809), whereas ensemble (random forest, gradient boosting) and linear or kernel methods performed similarly, indicating that discriminative information is consistently present across algorithmic families.

In two-dimensional UMAP projections, normal samples formed local groups but were not globally separated from tumors, either with all 2,863 features or with the 876 features that passed the Mann–Whitney U test with FDR correction when applied to all samples (for visualization only; silhouette −0.01 and 0.03; Fig 4). Hence, the tumor–normal signal is distributed across many fluxes and is captured by multivariate classifiers rather than by a low-dimensional projection.

**Fig 4.**
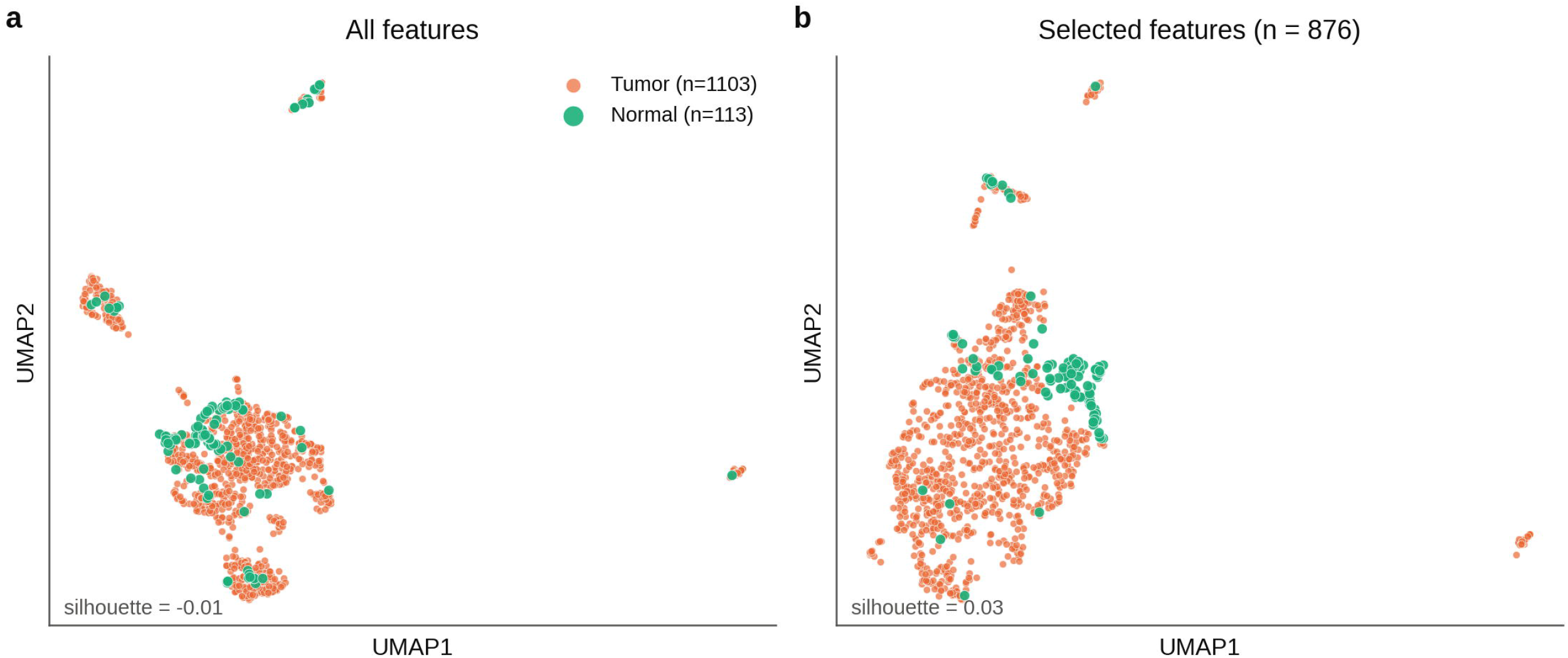
UMAP plot of selected features. Two-dimensional UMAP projection of tumor and normal samples using all features (a) and the 876 features selected by the Mann–Whitney U test with FDR correction applied to all samples, for visualization only (b); the classifiers selected features within each training fold. The axes of the two panels are not comparable because each embedding was computed independently.

Confusion matrices and ROC curves of the out-of-fold predictions confirmed robust classification across models (Figs 5 and 6), and evaluation on the independent, patient-grouped test set gave results consistent with cross-validation (Table 3).

**Fig 5.**
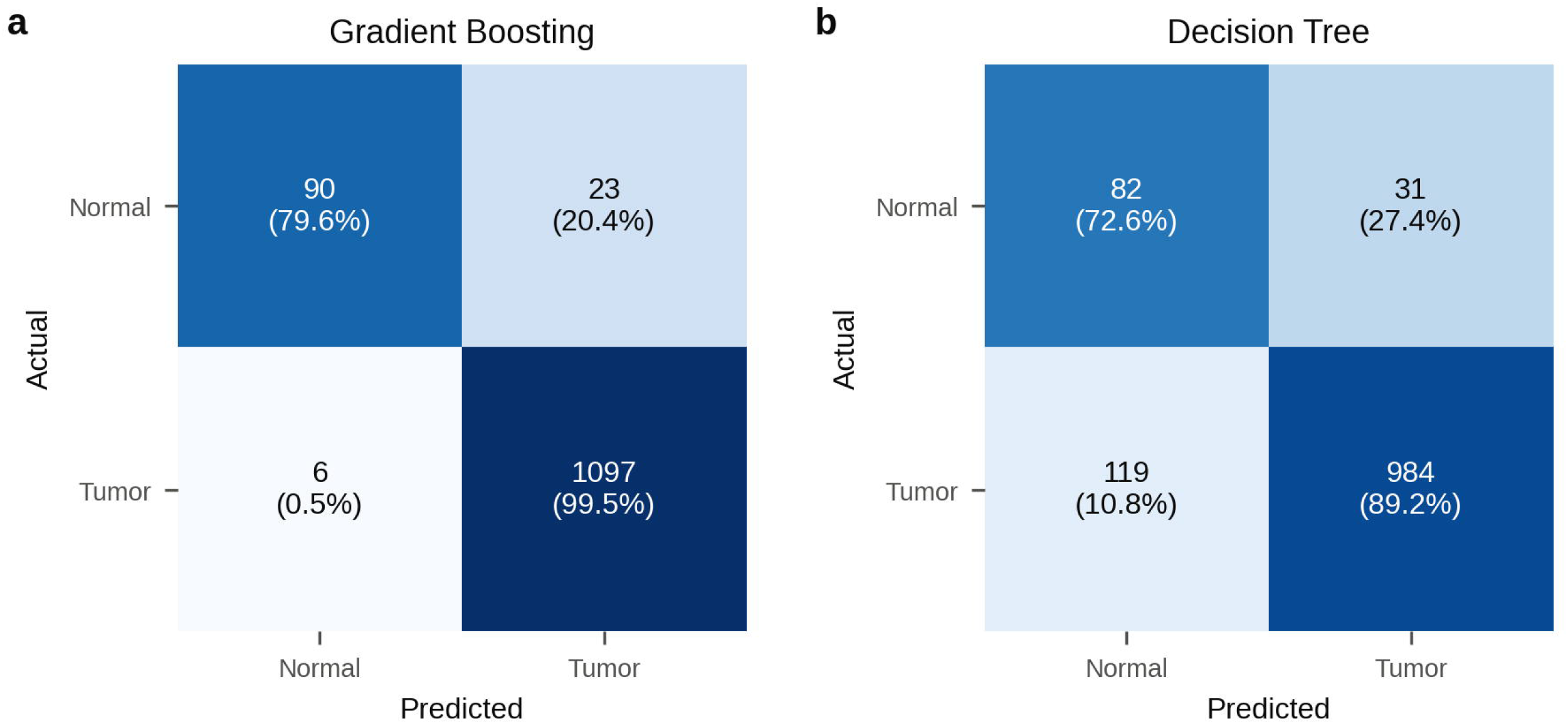
Confusion matrices of the Gradient Boosting and Decision Tree algorithms. Out-of-fold predictions of the nested cross-validation for the best (a) and the least accurate tree-based (b) classifier; values are counts and row percentages.

**Fig 6.**
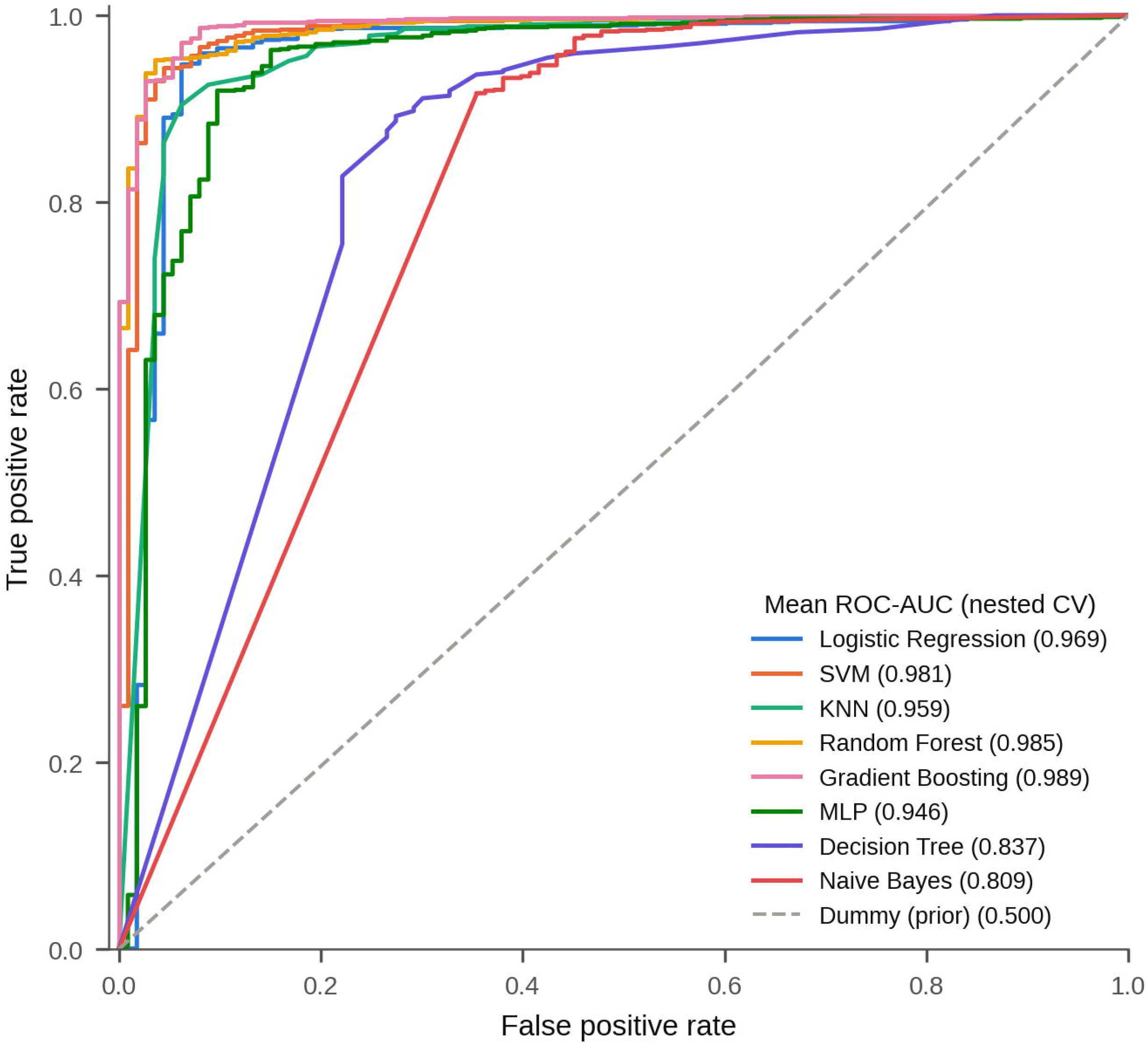
ROC curves. Out-of-fold ROC curves of the eight algorithms evaluated by nested cross-validation. The area under the curve ranges from 0.5 (random performance) to 1.0 (maximum performance).

**Fig 7.**
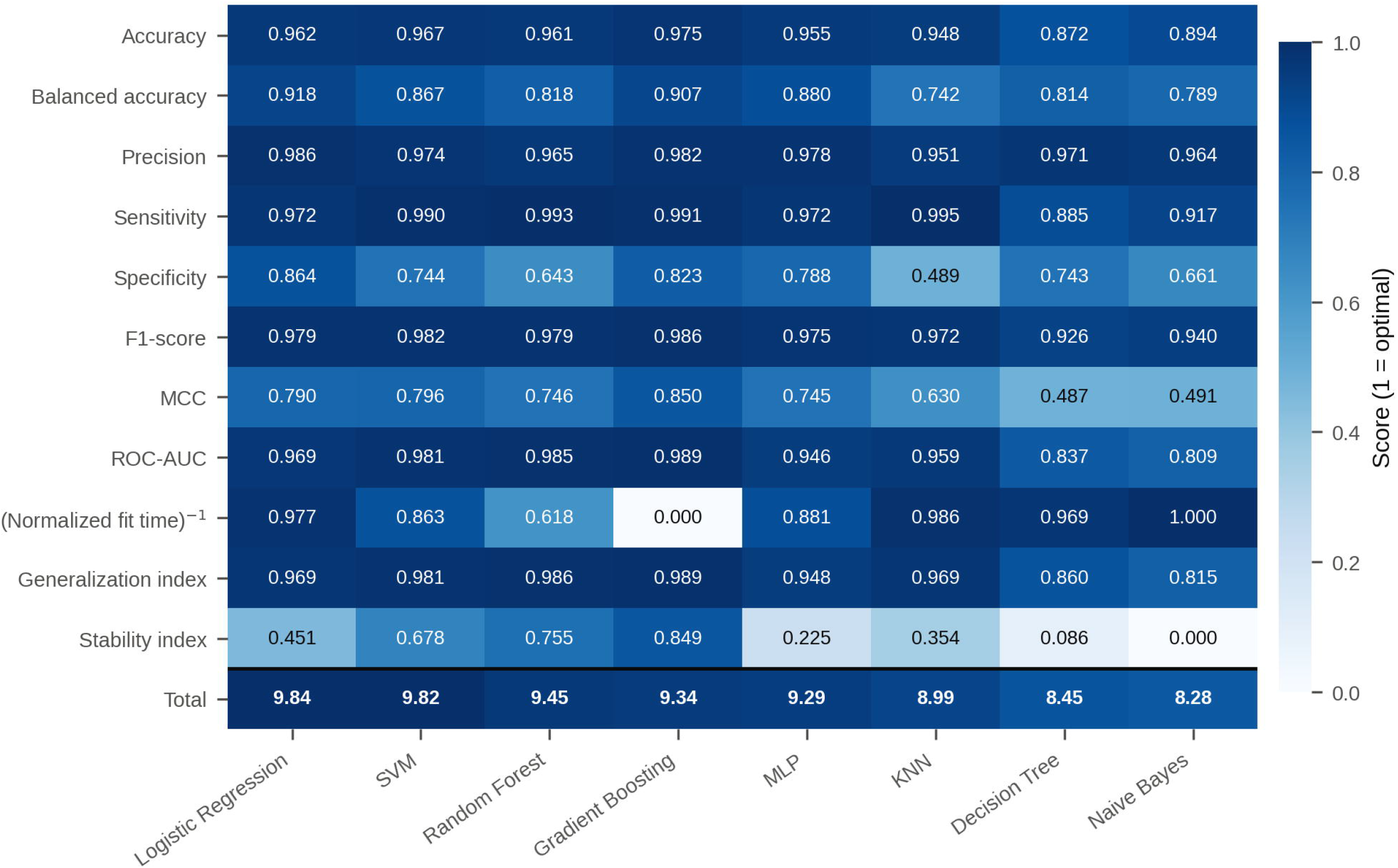
Decision matrix. Comparison of algorithm performance (columns) using various evaluation metrics (rows), including a total score. Values are normalized on a scale from 0 to 1 (1 = optimal performance). The color scale indicates the proximity of each value to 1.

**Table 3.** Performance on the independent test set. Patient-grouped, stratified 30% hold-out (364 samples, 331 tumors) not used for model selection.

| **Algorithm** | **Accuracy** | **Balanced accuracy** | **MCC** | **ROC-AUC** |
|---|---|---|---|---|
| Gradient Boosting | 0.975 | 0.891 | 0.842 | 0.990 |
| Random Forest | 0.970 | 0.888 | 0.810 | 0.978 |
| SVM | 0.956 | 0.812 | 0.709 | 0.966 |
| Logistic Regression | 0.959 | 0.855 | 0.740 | 0.968 |
| KNN | 0.945 | 0.820 | 0.658 | 0.937 |
| MLP | 0.956 | 0.867 | 0.733 | 0.928 |
| Decision Tree | 0.890 | 0.899 | 0.591 | 0.925 |
| Naive Bayes | 0.879 | 0.824 | 0.499 | 0.871 |
| Dummy (prior) | 0.909 | 0.500 | 0.000 | 0.500 |

Thus, Gradient Boosting achieved the highest discrimination, whereas a multi-criteria evaluation that also considered generalization, stability and computational cost ranked Logistic Regression first, followed by SVM (Fig 7): Logistic Regression reached a comparable ROC-AUC (0.969) with the highest specificity (0.864) and balanced accuracy (0.918) and the lowest computational cost. A simple linear model on selected fluxes is therefore sufficient to recover the tumor–normal signal.

Beyond predictive performance, the similar results of linear, kernel and ensemble methods indicate that tumor and normal metabolic states are largely separable in the full flux space, without requiring highly non-linear decision boundaries.

Together, these results indicate that metabolic fluxes contain high-dimensional features capable of discriminating tumor states in this dataset.

### Metabolic fluxes reveal intrinsic tumor heterogeneity

Following validation of the metabolic signal, unsupervised learning was applied to explore intratumoral heterogeneity.

Dimensionality reduction using UMAP showed higher cluster separation than PCA for both metabolic (silhouette 0.83 vs 0.40) and clinical data (0.71 vs 0.42) (Table 4). However, silhouette values computed in the original feature space were low (≤ 0.21), indicating that the embeddings sharpen a structure that is weak in the original feature space.

**Table 4.** Silhouette scores for cluster separation using UMAP vs PCA, computed on clinical and metabolic data. Values for the best partition of each embedding; values in parentheses are computed in the original feature space.

| **Data Type** | **UMAP** | **PCA** |
|---|---|---|
| Clinical Data | 0.71 (0.12) | 0.42 (0.14) |
| Metabolic Data | 0.83 (0.21) | 0.40 (0.09) |

Agreement between clustering solutions was low: across the distinct metabolic and clinical partitions, the maximum adjusted Rand index was 0.018 (median −0.006; Fig 8), and the selected metabolic and clinical partitions did not agree beyond chance (ARI = 0.003, permutation p = 0.20; n = 1,069). The clinical partition agreed with PAM50 (ARI = 0.109) only through the receptor variables (ARI = 0.000 without them), and the metabolic partition agreed little (ARI = 0.048).

**Fig 8.**
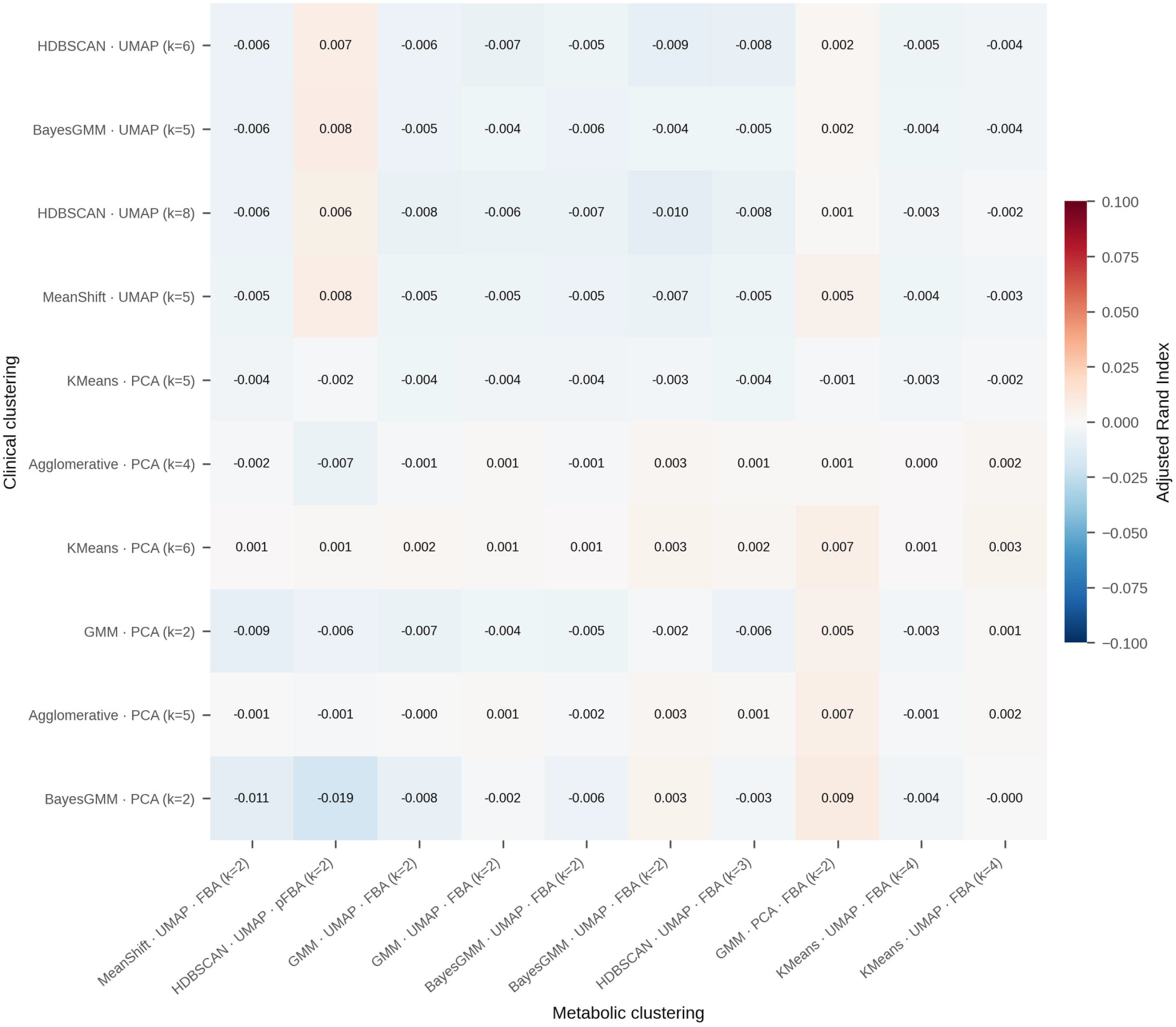
Concordance between clustering solutions. Adjusted Rand index (ARI) between the distinct partitions obtained from clinical data (rows) and from metabolic data (columns).

Of 366 candidate partitions, 280 were associated with medium supplementation and discarded. The selected partition (HDBSCAN on the UMAP embedding of pFBA metrics; stability across seeds, ARI = 0.97) separated 185 tumors (17%) from the remaining 871, with 36 tumors unassigned (Fig 10); the clinical partition identified three groups (Fig 9).

**Fig 9.**
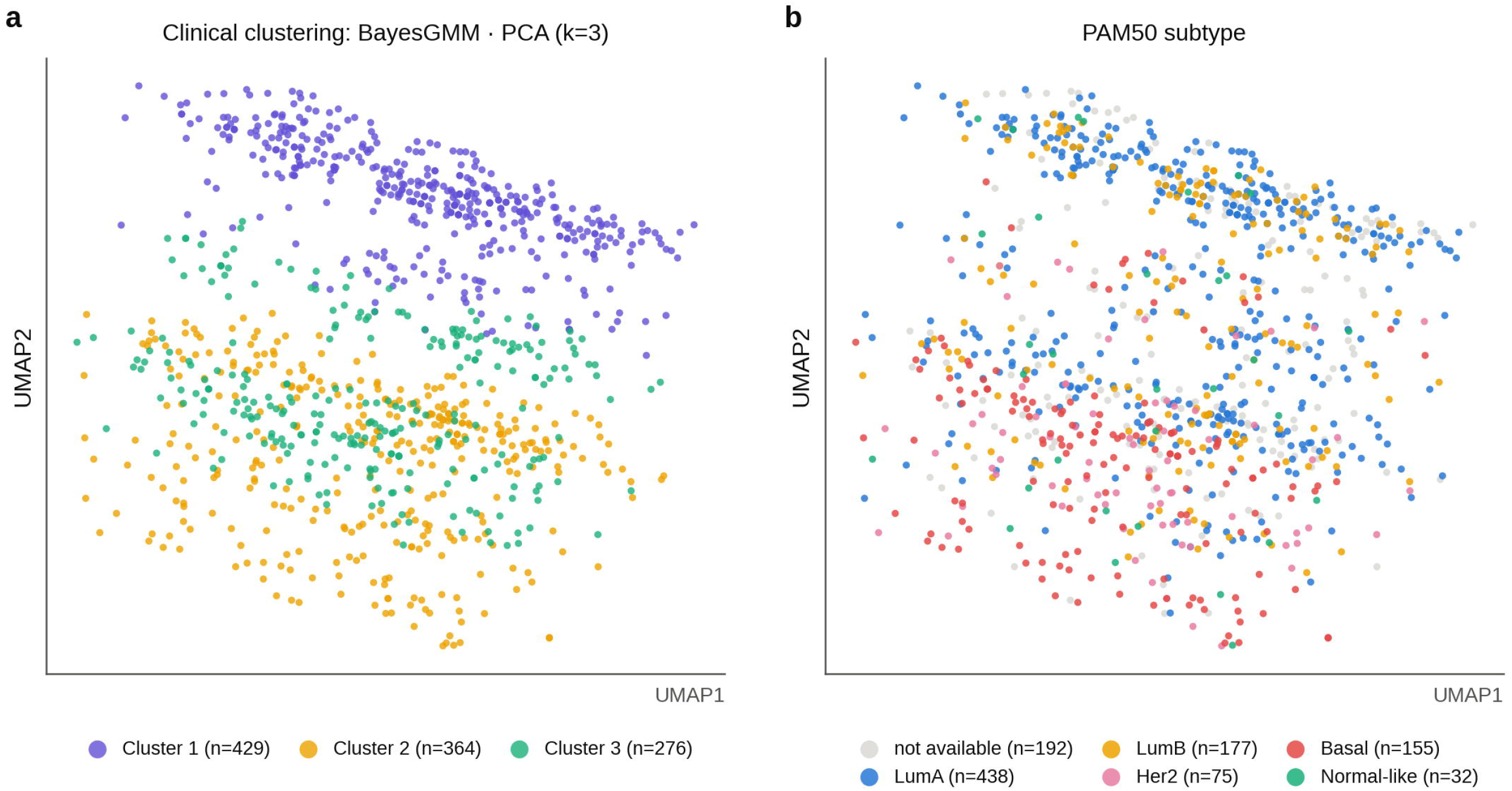
Clinical patient clustering. Two-dimensional UMAP embedding of the clinical variables; each point is a tumor, colored by clinical cluster (a) and by PAM50 subtype (b).

**Fig 10.**
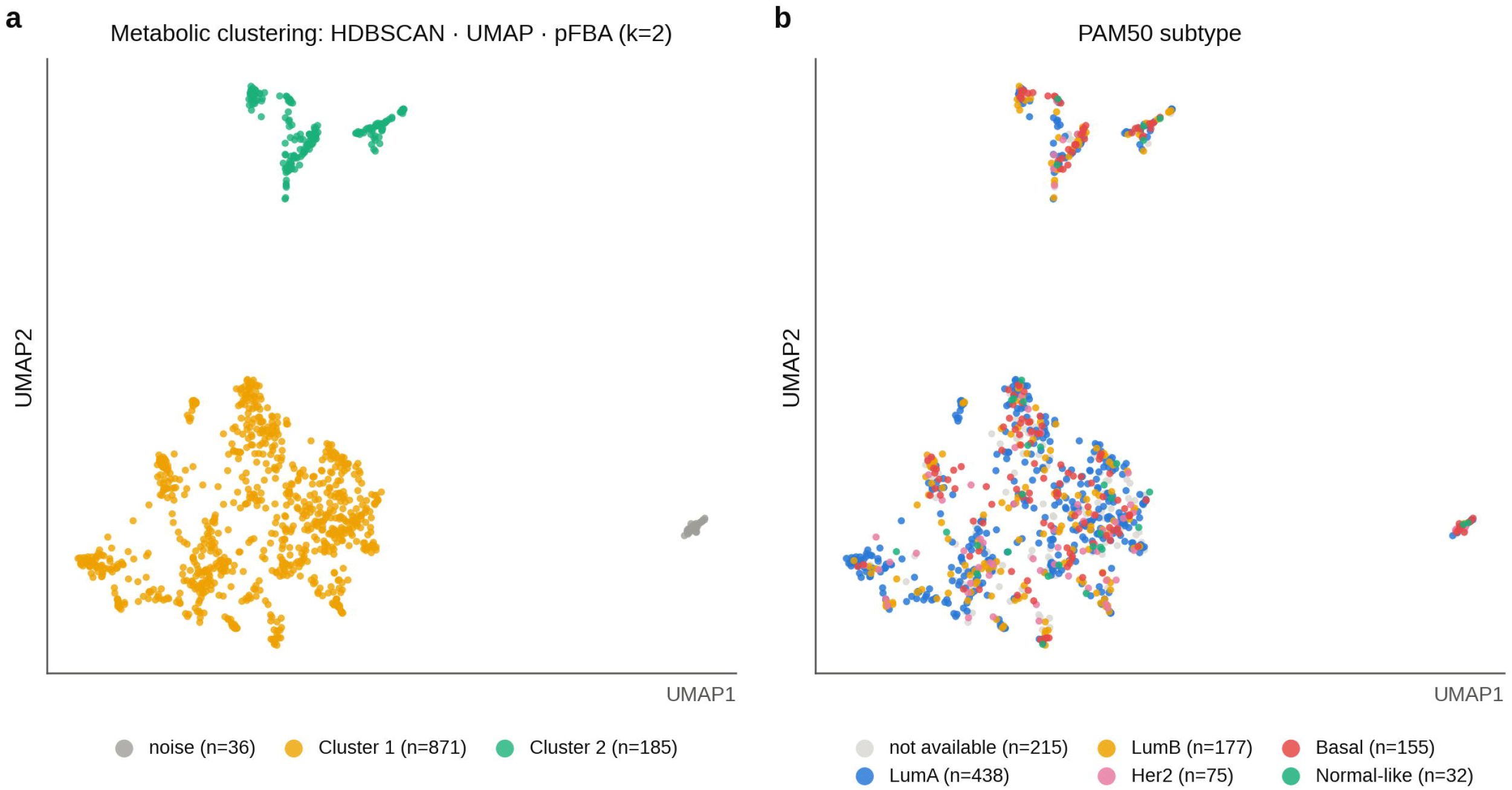
Metabolic patient clustering. Two-dimensional UMAP embedding of the pFBA metrics; each point is a tumor, colored by metabolic cluster (a; gray, unassigned) and by PAM50 subtype (b).

Clinical clusters were partly organized by receptor status, whereas both metabolic clusters contained tumors of all PAM50 subtypes (Figs 9 and 10).

Together, these observations indicate that the tumor metabolic space is heterogeneous but continuous rather than cleanly partitioned into discrete groups, and that it only partially overlaps with clinical classifications.

### Features of the metabolic clusters

The smaller cluster differed from the remaining tumors in 140 flux-derived features (Mann–Whitney U test, FDR ≤ 0.05), 66 with a large effect (|Cliff’s delta| ≥ 0.474; Fig 11; S6 Table). In both pFBA and L1w solutions, it showed lower biomass flux, carbon uptake, exchange and transport, and lower glycolytic, citric acid cycle and oxidative phosphorylation fluxes, together with higher enzyme allocation, consistent with a state of lower predicted growth. Its models were also of different size (p = 0.0005), so part of this difference may reflect reconstruction content. Using FBA metrics alone, the best partition instead isolated 36 tumors defined by phosphatidylinositol fluxes present only in the FBA solution; because FBA optima are not unique, it was not interpreted (ARI with the selected partition = 0.030).

**Fig 11.**
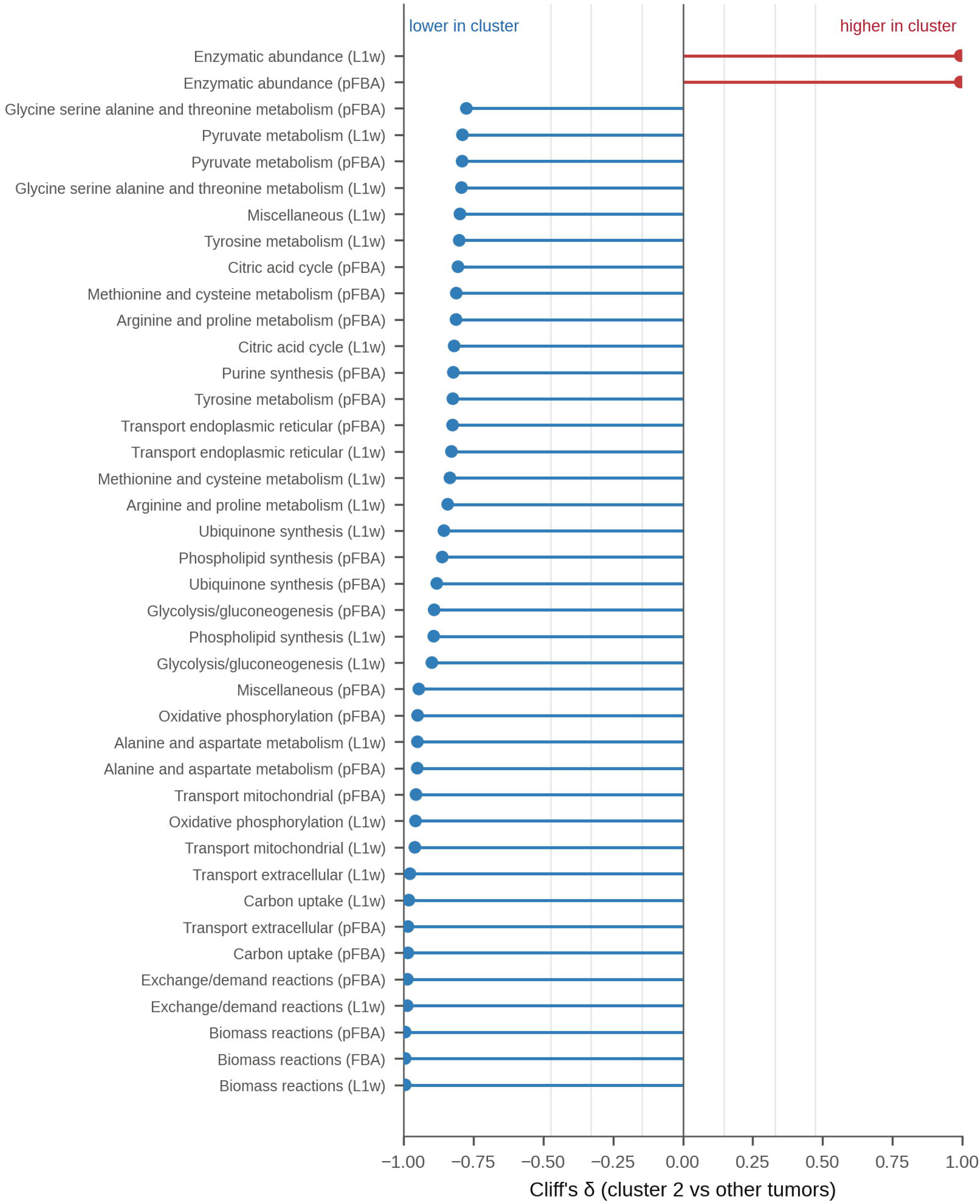
Cliff’s delta effect size analysis. Flux-derived features that differ significantly (FDR ≤ 0.05; at most 40 shown) ranked by Cliff’s delta between metabolic cluster 2 and the other tumors. Positive values indicate higher values in cluster 2; vertical lines mark the thresholds for small, medium and large effects (|δ| = 0.147, 0.33 and 0.474).

### Clinical characteristics predicted from metabolic fluxes

The metabolic clusters were not significantly associated with any clinical variable after FDR correction (strongest: ER, Cramér’s V = 0.063, FDR = 0.47; S6 Table). However, clustering is only one way of interrogating the flux space, and a continuous landscape need not split into discrete clusters even when it is biologically structured. When each clinical variable was predicted directly from reaction fluxes (Fig 12), fluxes predicted PAM50 Basal-like versus other subtypes (AUC = 0.966), triple-negative status (0.881), estrogen (0.863) and progesterone (0.770) receptor status, and Luminal A versus Luminal B (0.725), whereas stage (0.539), nodal status (0.506), age (0.545) and menopausal status (0.520) could not be predicted. HER2 status by immunohistochemistry was not predicted (0.513), and the HER2-enriched PAM50 subtype only weakly (0.624).

**Fig 12.**
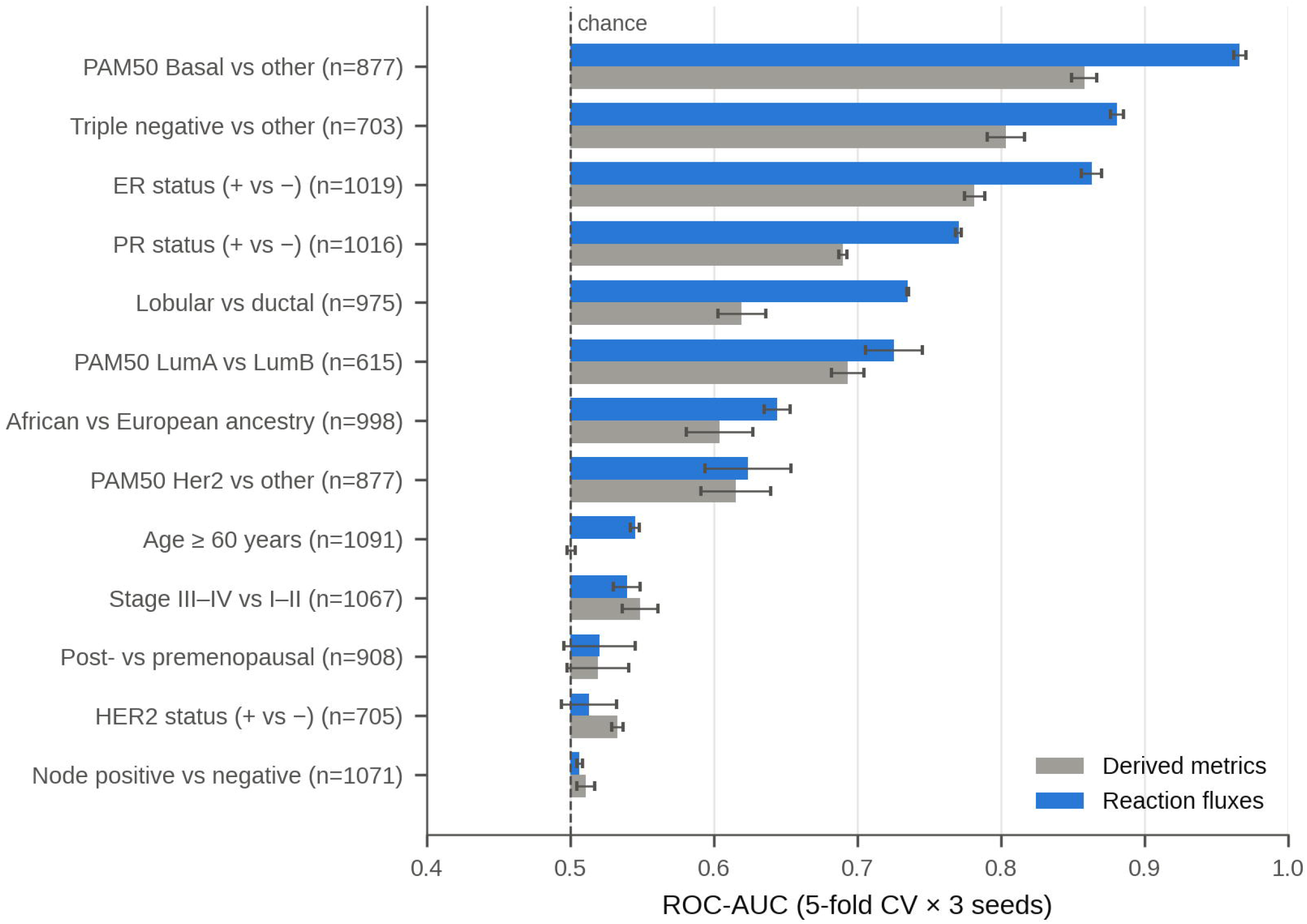
Clinical variables predicted from metabolic fluxes. ROC-AUC (mean ± SD over three repetitions of stratified 5-fold cross-validation) of a regularized logistic regression predicting each clinical variable from reaction fluxes or from derived metrics; n is the number of tumors with the variable available; the dashed line marks chance level (AUC = 0.5).

Overall survival was higher in the smaller metabolic cluster (log-rank p = 0.034; S2 Fig), but not after adjustment for age, stage and PAM50 subtype (HR = 0.63, 95% CI 0.35–1.14; 814 tumors, 107 deaths); survival results are therefore reported as exploratory in the Supporting Information.

Together with the low concordance between clustering solutions (Fig 8), these results suggest that metabolic states capture information that overlaps with, but is not fully explained by, clinical classifications.

### Metabolic signatures of breast cancer subtypes

Because receptor status and subtype were the clinical variables best predicted from fluxes, we characterized the metabolic differences among PAM50 subtypes. Of 212 flux-derived features (82 subsystem activities and derived metrics, each tested in the FBA, pFBA and L1w solutions in which it was available), 94 differed among subtypes (Kruskal–Wallis, FDR ≤ 0.05; Fig 13; S7 Table). These correspond to 52 distinct subsystems or metrics, 32 of which were significant in at least two solutions.

**Fig 13.**
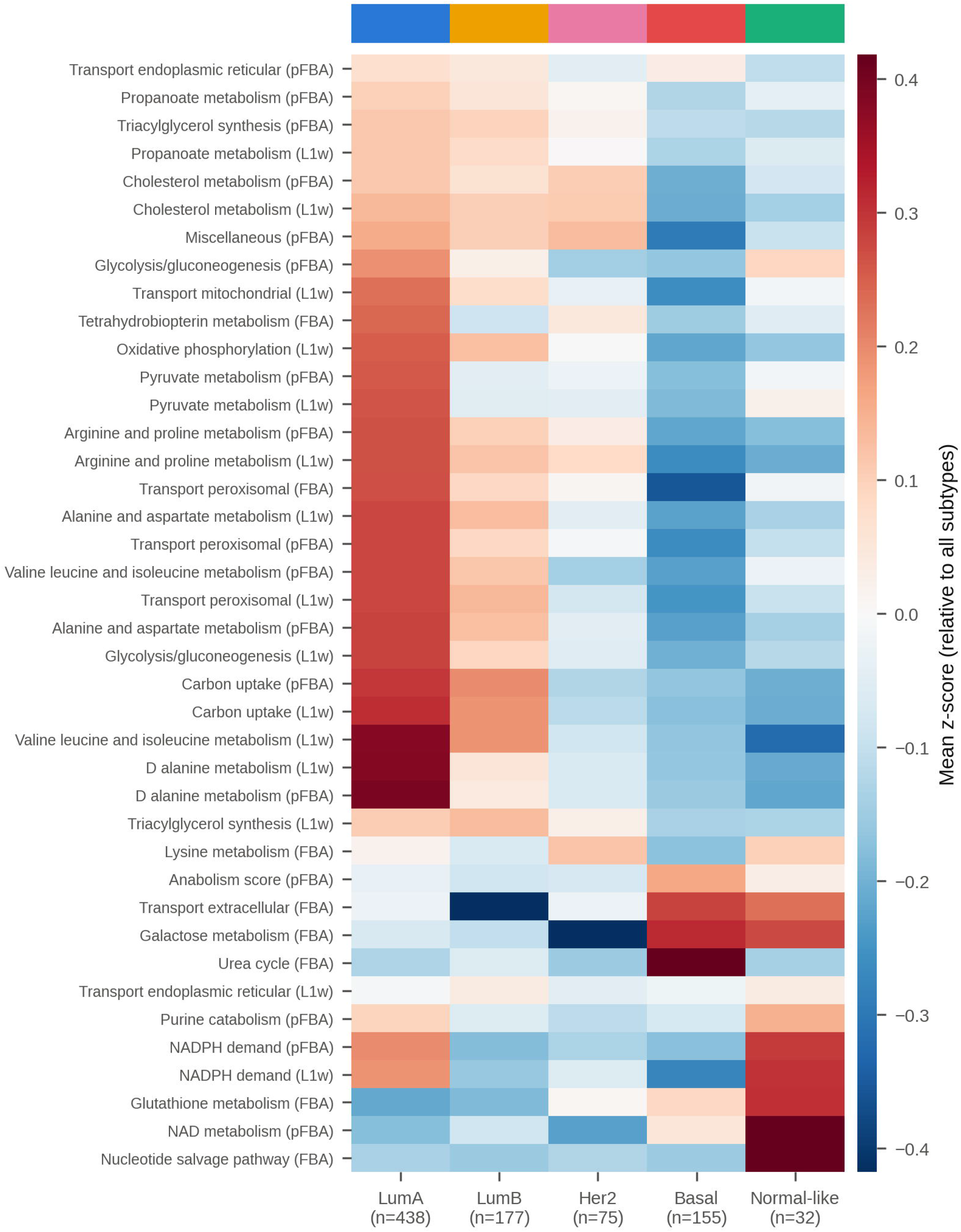
Metabolic signatures across molecular subtypes. Mean standardized activity (z-score, centered across subtypes) of the subsystem-level flux features that differ most among PAM50 subtypes (Kruskal–Wallis, FDR ≤ 0.05).

Luminal A models showed the highest activity in glycolysis, pyruvate metabolism, oxidative phosphorylation, carbon uptake, amino acid metabolism (branched-chain amino acids, alanine and aspartate, arginine and proline) and cholesterol and triacylglycerol synthesis. Basal-like models showed the lowest activity in most of these pathways but the highest activity in the urea cycle, extracellular transport, galactose metabolism and anabolism score. Luminal B tumors had profiles intermediate between Luminal A and the receptor-negative subtypes, and normal-like tumors showed higher NADPH demand and NAD, glutathione and nucleotide salvage activity (Fig 13).

These findings indicate that the metabolic organization captured by fluxes is largely aligned with established molecular classifications such as Luminal A/B or Basal-like, while the variation within subtypes may represent an additional, graded dimension of tumor heterogeneity.

## Discussion

### Metabolic fluxes capture structured functional information

Breast tissue is composed of multiple cell types that constitute the mammary gland, each exhibiting distinct metabolic characteristics. For example, basal cells tend to express abundant glycolytic enzymes, whereas luminal progenitors show increased levels of electron transport chain components, reflecting differences in energy metabolism among breast epithelial cell populations [35]. These differences indicate that metabolism varies across cell types within normal breast tissue.

During breast cancer development, alterations in transcription factors and signaling pathways such as MYC and TP53 drive proliferation and increase the demand for energy and biosynthetic precursors. Tumor cells therefore often take up more glucose and sustain glycolysis even when oxygen is available (the Warburg effect) [35].

Importantly, these metabolic alterations are not uniform across breast tumors and may vary according to tumor subtype, genetic alterations, and cellular composition [36]. Such metabolic heterogeneity can result in distinct patterns of energy production and biosynthetic activity among patients [36]. Therefore, characterizing metabolic states at the patient level may provide additional information for distinguishing tumors with different metabolic and proliferative phenotypes.

In this context, our analyses spanning reaction-level variability, classification and patient stratification showed that flux-derived features consistently captured variation associated with tumor state and heterogeneity. These representations capture reproducible differences between normal and tumor tissues, as well as among breast cancer subtypes, indicating that metabolic fluxes are associated with coordinated functional states rather than isolated pathway alterations.

Taken together, these findings suggest that tumor metabolism is structured mainly by receptor status and molecular subtype.

### Flux-based representations enable discrimination and reveal data structure

The consistently high performance across algorithms, with Gradient Boosting and Random Forest reaching the highest ROC-AUC under nested, patient-grouped cross-validation, shows that flux profiles contain robust discriminatory information, and the comparison between model classes indicates how this information is structured.

The similar performance of linear and ensemble methods suggests that tumor and normal samples are separable along many small, coordinated flux differences rather than a few dominant reactions. Consistent with previous studies integrating machine learning with genome-scale metabolic models [16–18,37], these findings suggest that metabolic states are embedded in a structured yet complex feature space, reflecting non-trivial dependencies across the metabolic network [38,39].

These results should not be interpreted as evidence that one learning paradigm is inherently superior. Rather, supervised approaches reveal strong discriminative signals between predefined states, while unsupervised analyses and the direct prediction of clinical variables show that flux distributions also contain structure associated with receptor status and molecular subtype. The combination of both perspectives supports the view that metabolic fluxes capture both separable and internally organized biological variation.

While classification performance alone does not establish mechanistic causality, it indicates that flux-derived features capture patterns consistently associated with tumor states.

### Metabolic heterogeneity emerges as a structured property

Our results suggest that breast cancer metabolism may be represented as a structured but continuous landscape rather than a small number of discrete metabolic states. Despite a conserved metabolic backbone, patient-specific models occupy different regions of the space of feasible metabolic configurations. Part of this variation is shared by tumors and distinguishes them from normal tissue; part follows receptor status and molecular subtype; and part is not explained by clinical classifications, as shown by the low agreement between metabolic and clinical partitions and by the weak cluster structure in the original feature space [4,9]. Metabolic states therefore capture information that overlaps with, but is not fully explained by, clinical classifications, suggesting that patient-specific fluxes can provide additional information for tumor characterization.

### Subtype-specific metabolism in breast cancer

Several of the metabolic differences among subtypes agree with experimental observations in breast tumors. Triple-negative and Basal-like tumors depend strongly on the uptake of extracellular nutrients such as glutamine and cystine, the latter through the xCT antiporter [40,41], consistent with the higher extracellular transport activity inferred here for Basal-like models. In contrast, Basal-like tumors express higher levels of glycolytic enzymes and glucose transporters than luminal tumors [42], whereas our models assigned the highest glycolytic flux to Luminal A tumors. Flux balance solutions favor routes with high ATP and biomass yield and do not include enzyme capacity constraints, which limits their ability to reproduce aerobic glycolysis, so glycolytic fluxes should be interpreted with caution.

Luminal tumors, in contrast, rely more on oxidative and lipid metabolism, including hormone-regulated lipogenesis [43,44], in line with the higher oxidative phosphorylation, cholesterol and triacylglycerol synthesis fluxes of Luminal A models.

The fact that receptor status and subtype, but not stage, nodal status or age, were recoverable from fluxes suggests that the models capture metabolic programs associated with tumor lineage rather than disease extent. Because each subtype contains tumors with heterogeneous fluxes, metabolic variation within subtypes may still carry information relevant to therapy, as suggested by pathway-based subtyping of triple-negative tumors [8].

From a modeling perspective, these subtype differences are associated with different regions of the feasible flux space under a common medium, although this interpretation depends on model assumptions and requires experimental validation.

### Clinical relevance and translational implications

The association between flux-derived features and receptor status and subtype suggests that metabolic models translate transcriptomic profiles into functional states that are consistent with clinical classifications, while metabolic clusters were not independently associated with survival.

From a therapeutic perspective, subtype-specific dependencies, such as glutamine and cystine uptake in triple-negative tumors and lipogenesis in luminal and HER2-positive tumors, are candidate metabolic vulnerabilities [40,41,43], and patient-specific models may help to identify which tumors within a subtype share them. Associations between metabolic phenotypes and clinical outcomes have been reported across cancer types [45,46], but larger cohorts with treatment information would be required to evaluate prognostic value.

An important translational observation is that clinical variables alone produced groupings that were more closely related to molecular subtype than metabolic clusters, whereas the information shared by both is captured by receptor status and subtype. This suggests that routinely collected clinical data and flux-based models provide complementary rather than redundant descriptions of tumors.

More generally, flux inference transforms transcriptomic data into predictions of cellular activity: transcript abundance does not indicate which reactions are active or at what rates, but embedding it in a network of stoichiometric and physicochemical constraints limits the possible states to those that are biochemically feasible and consistent with expression.

Flux estimation can thus be viewed as a biologically informed filter that retains only metabolically consistent configurations, which may help to identify coordinated functional patterns that are not apparent from gene expression alone.

This perspective is consistent with large-scale efforts such as TCGA, which provide extensive molecular data but are inherently limited in their ability to capture cellular function [22]. Integrating these data into mechanistic frameworks may therefore enhance their interpretability by linking molecular measurements to functional phenotypes.

### Positioning within the current literature

This work builds on previous studies integrating genome-scale metabolic modeling with machine learning [14,17,18], but differs in its focus on the intrinsic structure of flux-derived features. Rather than treating metabolic outputs as auxiliary variables, the present results suggest that fluxes themselves capture functional organization.

In contrast to pathway-specific subtyping approaches [8], this framework captures coordinated changes across the metabolic network, enabling the characterization of system-level differences among subtypes and complementing ongoing efforts in precision medicine [24].

### Limitations and methodological considerations

Several limitations should be considered. Fluxes are inferred from constraint-based models rather than measured, and depend on modeling choices: the objective, the expression threshold for active genes (tested here at a single value), the curated constraints, and the simulated medium; actual nutrient availability varies between and within tumors. FBA optima are not unique, so partitions of tumors did not rely on FBA alone. Bulk RNA-seq mixes tumor, stromal and immune cells, so the models reflect tissue-level metabolism, and tumor purity was not modeled. Some reconstructions contain internal cycles that regenerate ATP without uptake, which may inflate ATP-related metrics. Finally, transcript levels only indirectly reflect enzyme activity, steady state ignores temporal and spatial heterogeneity, and whether fluxes add predictive value beyond expression-based classifiers was not tested.

Therefore, the results should be interpreted as computationally derived hypotheses requiring experimental validation. Additionally, model performance and generalizability may depend on cohort composition and should be evaluated in independent datasets.

### Implications for precision medicine

Despite these limitations, the association of flux-derived features with receptor status and molecular subtype highlights the potential of flux-based analysis as a complementary layer of tumor characterization, in particular for identifying subtype-specific metabolic dependencies.

More broadly, these results support a framework in which tumor metabolism constitutes an additional dimension of heterogeneity, linking molecular constraints to functional phenotypes and potentially improving the interpretability of patient-specific models.

## Conclusion

This study provides evidence that metabolic fluxes derived from genome-scale models capture information associated with tumor state, receptor status and molecular subtype, but not with disease extent. Together, these findings support a view of breast cancer metabolism as a structured but continuous landscape, in which patient-specific fluxes capture both a shared tumor metabolic state and patient-to-patient heterogeneity that relates to, but is not reduced to, conventional molecular and clinical classifications.

## Availability of Data and Materials

The code used in this study is available at https://github.com/GIMIudg/GIMIpapers/tree/main/Precision-Oncology-for-Breast-Cancer-Diagnosis and is archived in Zenodo [47]. The derived data (flux solutions, model quality-control indicators, and the tables and figures of the analysis) are archived in Zenodo [48]. The raw TCGA-BRCA expression, clinical and survival data are not redistributed; they are publicly available from the UCSC Xena GDC hub (https://xenabrowser.net).

## Acknowledgments

We thank the Secretaría de Ciencia, Humanidades, Tecnología e Innovación (SECIHTI) for funding this research (CVU No. 1347052).

## Supporting Information

**S1 File. Bibliomics data.** Excel file of bibliomic data (curated breast cancer–associated genes, reactions and metabolites) in a format compatible with XomicsToModel, with literature references.

**S1 Table. Selected clinical data. Clinical variables used for the clinical clustering.**

**S2 Table. Metabolic optimization description.** Metabolic optimization characteristics for unsupervised training data.

**S3 Table. Algorithm configurations.** Hyperparameter search spaces, selected hyperparameters, and per-fold performance (mean ± SD) of all classifiers; dimensionality reduction and clustering parameters.

**S4 Table. Uniform medium.** Exchange reactions open for uptake, HPLM concentrations, and uptake bounds applied to all models; per-model supplementation needed to reach the minimal biomass flux.

**S5 Table. Tumor versus normal reactions.** Reactions whose flux differs between tumor and normal samples, with subsystem, Cliff’s delta, FDR, and associated genes.

**S6 Table. Metabolic clusters.** Effect sizes of flux-derived features between metabolic clusters and association of clusters with clinical variables.

**S7 Table. PAM50 metabolic features.** Kruskal–Wallis tests of flux-derived features among PAM50 subtypes.

**S1 Fig. Comparison of flux solutions.** Number of reactions carrying flux (a), tumor–normal ROC-AUC (b) and median per-model correlation between solutions (c) for the FBA, pFBA, L1w, L2 and L2w solutions.

**S2 Fig. Survival analysis.** Kaplan–Meier curves of overall survival by metabolic cluster (a) and by PAM50 subtype (b), with 95% confidence bands and log-rank p-values; the Cox model adjusted for age, stage and PAM50 subtype is given in S6 Table.

**S3 Fig. Multi-objective optimization.** Minimal carbon uptake (a) and enzyme allocation (b) along the biomass axis of the biomass–ATP grid for tumor and normal models (median and interquartile range).

## References

[1] Goetz LH, Schork NJ. Personalized medicine: motivation, challenges, and progress. Fertil Steril. 2018;109(6):952–963. doi: 10.1016/j.fertnstert.2018.05.006

[2] Pal S, Mondal S, Das G, Khatua S, Ghosh Z. Big data in biology: the hope and present-day challenges in it. Gene Rep. 2020;21:100869. doi: 10.1016/j.genrep.2020.100869

[3] Hasin Y, Seldin M, Lusis A. Multi-omics approaches to disease. Genome Biol. 2017;18:83. doi: 10.1186/s13059-017-1215-1

[4] Yizhak K, Chaneton B, Gottlieb E, Ruppin E. Modeling cancer metabolism on a genome scale. Mol Syst Biol. 2015;11(6):817. doi: 10.15252/msb.20145307

[5] Gu C, Kim GB, Kim WJ, Kim HU, Lee SY. Current status and applications of genome-scale metabolic models. Genome Biol. 2019;20:121. doi: 10.1186/s13059-019-1730-3

[6] Hanahan D, Weinberg RA. Hallmarks of cancer: The next generation. Cell. 2011;144(5):646–674. doi: 10.1016/j.cell.2011.02.013

[7] Guo L, Kong D, Liu J, Zhan L, Luo L, Zheng W, et al. Breast cancer heterogeneity and its implication in personalized precision therapy. Exp Hematol Oncol. 2023;12(1):3. doi: 10.1186/s40164-022-00363-1

[8] Gong Y, Ji P, Yang YS, Xie S, Yu TJ, Xiao Y, et al. Metabolic-pathway-based subtyping of triple-negative breast cancer reveals potential therapeutic targets. Cell Metab. 2021;33(1):51–64. doi: 10.1016/j.cmet.2020.10.012

[9] Richelle A, Chiang AWT, Kuo CC, Lewis NE. Increasing consensus of context-specific metabolic models by integrating data-inferred cell functions. PLoS Comput Biol. 2019;15(4):e1006867. doi: 10.1371/journal.pcbi.1006867

[10] Heirendt L, Arreckx S, Pfau T, Mendoza SN, Richelle A, Heinken A, et al. Creation and analysis of biochemical constraint-based models using the COBRA Toolbox v. 3.0. Nat Protoc. 2019;14(3):639–702. doi: 10.1038/s41596-018-0098-2

[11] Brunk E, Sahoo S, Zielinski DC, Altunkaya A, Dräger A, Mih N, et al. Recon3D enables a three-dimensional view of gene variation in human metabolism. Nat Biotechnol. 2018;36(3):272–281. doi: 10.1038/nbt.4072

[12] Ravi S, Gunawan R. ΔFBA—Predicting metabolic flux alterations using genome-scale metabolic models and differential transcriptomic data. PLoS Comput Biol. 2021;17(11):e1009589. doi: 10.1371/journal.pcbi.1009589

[13] Preciat G, Wegrzyn AB, Luo X, Thiele I, Hankemeier T, Fleming RMT, et al. XomicsToModel: omics data integration and generation of thermodynamically consistent metabolic models. Nat Protoc. 2025 Dec 1. doi: 10.1038/s41596-025-01288-9

[14] Magazzù G, Zampieri G, Angione C. Clinical stratification improves the diagnostic accuracy of small omics datasets within machine learning and genome-scale metabolic modelling methods. Comput Biol Med. 2022;151:106244. doi: 10.1016/j.compbiomed.2022.106244

[15] MacEachern SJ, Forkert ND. Machine learning for precision medicine. Genome. 2021;64(4):416–425. doi: 10.1139/gen-2020-0131

[16] Antonakoudis A, Barbosa R, Kotidis P, Kontoravdi C. The era of big data: Genome-scale modelling meets machine learning. Comput Struct Biotechnol J. 2020;18:3287–3300. doi: 10.1016/j.csbj.2020.10.011

[17] Lewis JE, Kemp ML. Integration of machine learning and genome-scale metabolic modeling identifies multi-omics biomarkers for radiation resistance. Nat Commun. 2021;12:2700. doi: 10.1038/s41467-021-22989-1

[18] Lee SM, Lee GR, Kim HU. Machine learning-guided evaluation of extraction and simulation methods for cancer patient-specific metabolic models. Comput Struct Biotechnol J. 2022;20:3041–3052. doi: 10.1016/j.csbj.2022.06.027

[19] Lu J, Liu P, Zhang R. A metabolic gene signature to predict breast cancer prognosis. Front Mol Biosci. 2022;9:900433. doi: 10.3389/fmolb.2022.900433

[20] Prusty S, Patnaik S, Dash SK. SKCV: Stratified K-fold cross-validation on ML classifiers for predicting cervical cancer. Front Nanotechnol. 2022;4:972421. doi: 10.3389/fnano.2022.972421

[21] Saeys Y, Inza I, Larranaga P. A review of feature selection techniques in bioinformatics. Bioinformatics. 2007;23(19):2507–2517. doi: 10.1093/bioinformatics/btm344

[22] Tomczak K, Czerwińska P, Wiznerowicz M. The Cancer Genome Atlas (TCGA): an immeasurable source of knowledge. Contemp Oncol. 2015;19(1):68–77. doi: 10.5114/wo.2014.47136

[23] Baloni P, Dinalankara W, Earls JC, Knijnenburg TA, Geman D, Marchionni L, et al. Identifying personalized metabolic signatures in breast cancer. Metabolites. 2020;11(1):20. doi: 10.3390/metabo11010020

[24] Turanli B, Gulfidan G, Aydogan OO, Kula C, Selvaraj G, Arga KY. Genome-scale metabolic models in translational medicine: the current status and potential of machine learning in improving the effectiveness of the models. Mol Omics. 2024;20(4):234–247. doi: 10.1039/D3MO00152K

[25] Cantor JR, Abu-Remaileh M, Kanarek N, et al. Physiologic medium rewires cellular metabolism and reveals uric acid as an endogenous inhibitor of UMP synthase. Cell. 2017;169(2):258–272.e17. doi: 10.1016/j.cell.2017.03.023

[26] Agren R, Bordel S, Mardinoglu A, Pornputtapong N, Nookaew I, Nielsen J. Reconstruction of Genome-Scale Active Metabolic Networks for 69 Human Cell Types and 16 Cancer Types Using INIT. PLoS Comput Biol. 2012;8(5):e1002518. doi: 10.1371/journal.pcbi.1002518

[27] Lewis NE, Hixson KK, Conrad TM, et al. Omic data from evolved E. coli are consistent with computed optimal growth from genome-scale models. Mol Syst Biol. 2010;6:390. doi: 10.1038/msb.2010.47

[28] Machado D, Herrgård M. Systematic evaluation of methods for integration of transcriptomic data into constraint-based models of metabolism. PLoS Comput Biol. 2014;10(4):e1003580. doi: 10.1371/journal.pcbi.1003580

[29] Blais EM, Rawls KD, Dougherty BV, et al. Reconciled rat and human metabolic networks for comparative toxicogenomics and biomarker predictions. Nat Commun. 2017;8:14250. doi: 10.1038/ncomms14250

[30] Dai Z, Yang S, Xu L, et al. Identification of cancer-associated metabolic vulnerabilities by modeling multi-objective optimality in metabolism. Cell Commun Signal. 2019;17:124. doi: 10.1186/s12964-019-0439-y

[31] Fay MP, Proschan MA. Wilcoxon-Mann-Whitney or t-test? On assumptions for hypothesis tests and multiple interpretations of decision rules. Stat Surv. 2010;4:1–39. doi: 10.1214/09-SS051

[32] Ferreira JA. The Benjamini-Hochberg Method in the Case of Discrete Test Statistics. Int J Biostat. 2007;3(1). doi: 10.2202/1557-4679.1065

[33] Xu R, Wunsch D. Survey of clustering algorithms. IEEE Trans Neural Netw. 2005;16(3):645–678. doi: 10.1109/TNN.2005.845141

[34] Hubert L, Arabie P. Comparing partitions. J Classif. 1985;2:193–218. doi: 10.1007/BF01908075

[35] Wang CC. Metabolic stress adaptations underlie mammary gland morphogenesis and breast cancer progression. Cells. 2021;10(10):2641. doi: 10.3390/cells10102641

[36] Patra S, Elahi N, Armorer A, Arunachalam S, Omala J, Hamid I, et al. Mechanisms governing metabolic heterogeneity in breast cancer and other tumors. Front Oncol. 2021;11:700629. doi: 10.3389/fonc.2021.700629

[37] Tanıl E, Kızılilsoley N, Nikerel E. Genome-scale Metabolic Model guided subtyping lung cancer towards personalized diagnosis. IFAC Pap Online. 2022;55(20):641–646. doi: 10.1016/j.ifacol.2022.09.168

[38] Vander Heiden MG, Cantley LC, Thompson CB. Understanding the Warburg effect: The metabolic requirements of cell proliferation. Science. 2009;324(5930):1029–1033. doi: 10.1126/science.1160809

[39] Pavlova NN, Thompson CB. The emerging hallmarks of cancer metabolism. Cell Metab. 2016;23(1):27–47. doi: 10.1016/j.cmet.2015.12.006

[40] Kim S, Kim DH, Jung WH, Koo JS. Expression of glutamine metabolism-related proteins according to molecular subtype of breast cancer. Endocr Relat Cancer. 2013;20(3):339–348. doi: 10.1530/ERC-12-0398

[41] Timmerman LA, Holton T, Yuneva M, et al. Glutamine sensitivity analysis identifies the xCT antiporter as a common triple-negative breast tumor therapeutic target. Cancer Cell. 2013;24(4):450–465. doi: 10.1016/j.ccr.2013.08.020

[42] Choi J, Kim DH, Jung WH, Koo JS. Metabolic interaction between cancer cells and stromal cells according to breast cancer molecular subtype. Breast Cancer Res. 2013;15(5):R78. doi: 10.1186/bcr3472

[43] Menendez JA, Lupu R. Fatty acid synthase and the lipogenic phenotype in cancer pathogenesis. Nat Rev Cancer. 2007;7(10):763–777. doi: 10.1038/nrc2222

[44] Gandhi N, Das GM. Metabolic reprogramming in breast cancer and its therapeutic implications. Cells. 2019;8(2):89. doi: 10.3390/cells8020089

[45] Wei X, Michelakos T, He Q, Wang X, Chen Y, Kontos F, et al. Association of tumor cell metabolic subtype and immune response with the clinical course of hepatocellular carcinoma. Oncologist. 2023;28(11):e1031–e1042. doi: 10.1093/oncolo/oyad113

[46] Li R, Zhu J, Zhong WD, Jia Z. Comprehensive evaluation of machine learning models and gene expression signatures for prostate cancer prognosis using large population cohorts. Cancer Res. 2022;82(9):1832–1843. doi: 10.1158/0008-5472.CAN-21-3074

[47] GIMIudg. GIMIpapers: Precision-Oncology-for-Breast-Cancer-Diagnosis [software]. Zenodo; 2026. doi: 10.5281/zenodo.19339596

[48] Ruiz Robles E, Rincón-Ballesteros R, Chacón Méndez SA, Alvarez-Padilla FJ, Preciat G. Patient-specific metabolic fluxes reflect molecular subtype and heterogeneity in breast cancer: analysis results [dataset]. Zenodo; 2026. doi: 10.5281/zenodo.23779079

