## Supplementary figures and images for "Patient-specific metabolic fluxes reflect molecular subtype and heterogeneity in breast cancer"

### Supplemental Figure 1

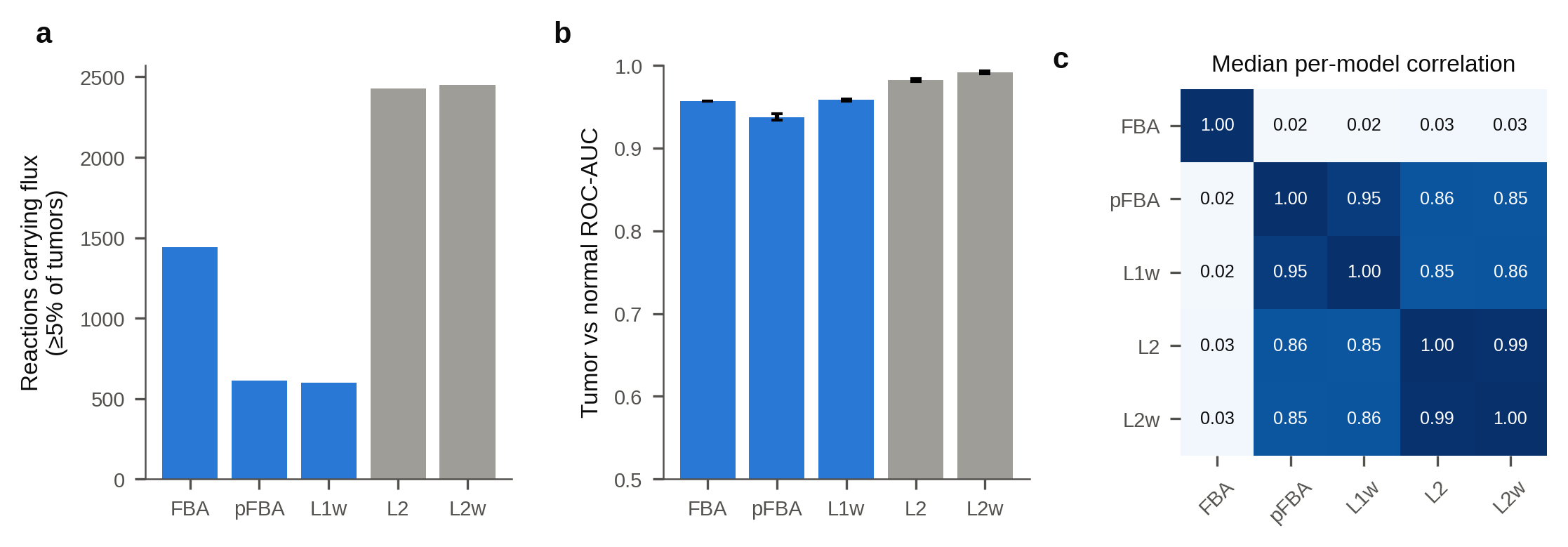

### Supplemental Figure 2

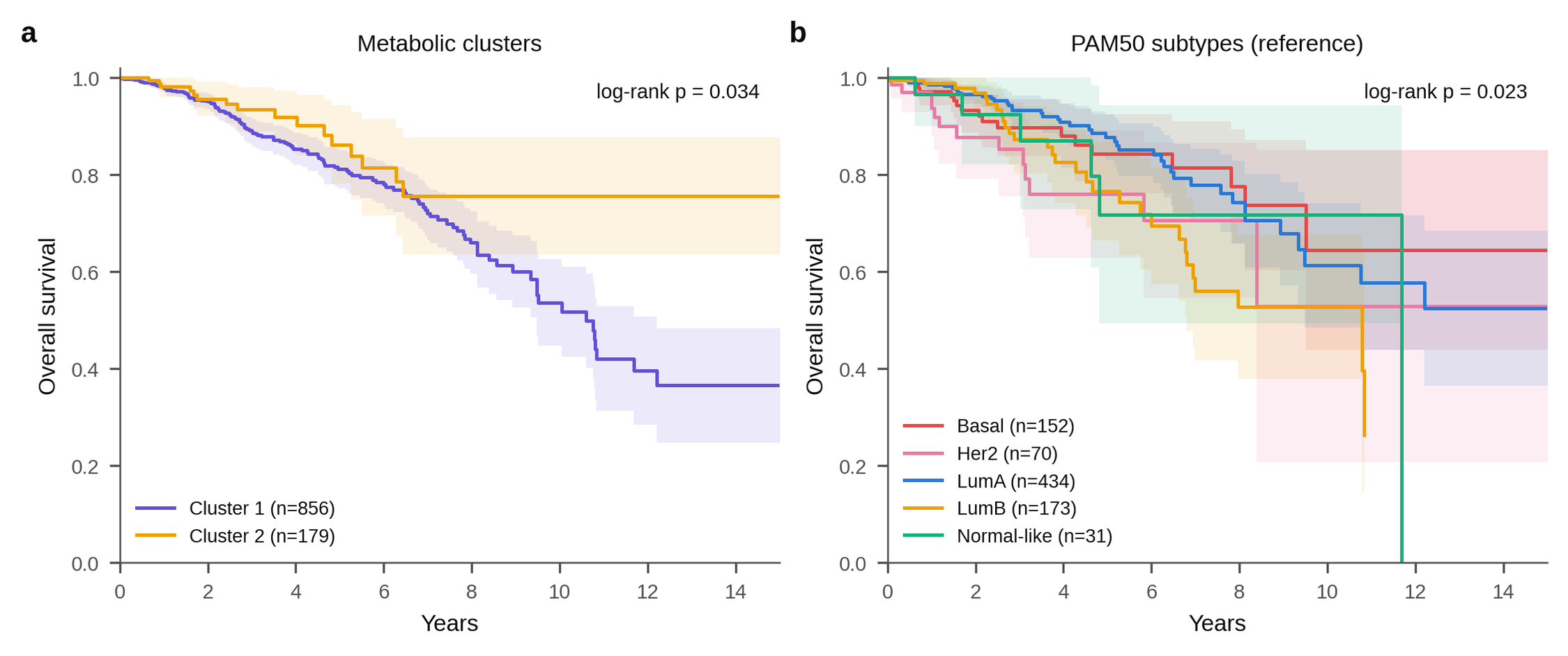

### Supplemental Figure 3

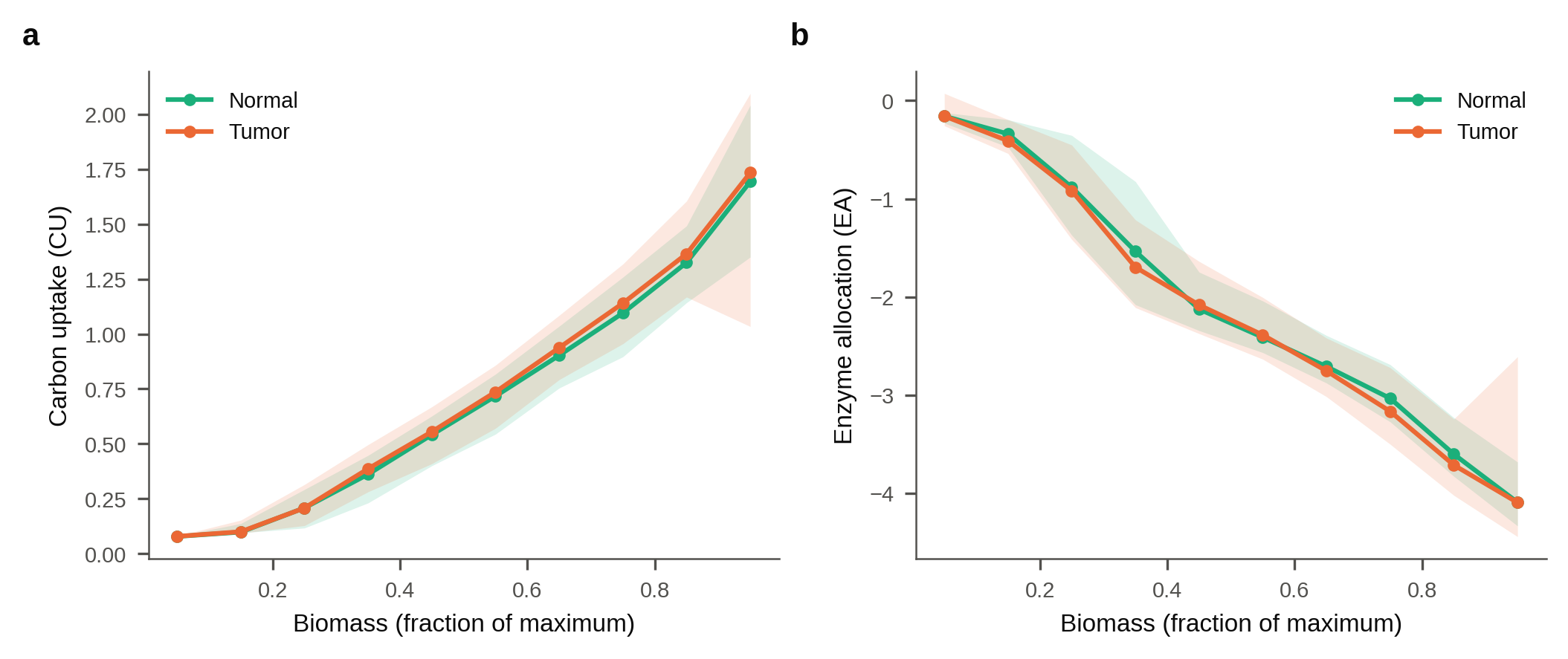
